# Interferon-γ lowers tumour growth by increasing glycolysis and lactate production in a nitric oxide-dependent manner: implications for cancer immunotherapy

**DOI:** 10.1101/2023.07.25.550474

**Authors:** Avik Chattopadhyay, Sirisha Jagdish, Aagosh Kishore Karhale, Nikita S. Ramteke, Arsha Zaib, Dipankar Nandi

## Abstract

Interferon-gamma (IFN-γ), the sole member of the type-II interferon family, is well known to protect the host from infectious diseases as well as mount anti-tumour responses. The amounts of IFN-γ in the tumour microenvironment determine the host responses against tumours; however, several tumours employ evasive strategies by responding to low IFN-γ signalling. In this study, the response of various tumour cell lines to IFN-γ was studied *in vitro*. IFN-γ-activation increases glycolytic flux and reduces mitochondrial function in a nitric oxide (NO)- and reactive oxygen species (ROS)-dependent manner in the H6 hepatoma tumour cell line. The higher glycolysis further fuelled NO and ROS production, indicating a reciprocal regulation. These processes are accompanied by Hypoxia inducing factor (HIF)-1*α* stabilization and HIF-1α-dependent augmentation of the glycolytic flux. The IFN-γ enhancement of lactate production also occurred in other NO-producing cell lines: RAW 264.7 monocyte/macrophage and Renca renal adenocarcinoma. However, two other tumour cell lines, CT26 colon carcinoma and B16F10 melanoma, did not produce NO and lactate upon IFN-γ-activation. HIF-1α stabilization upon IFN-γ-activation led to lower cell growth of B16F10 but not CT26 cells. Importantly, the IFN-γ-activation of both CT26 and B16F10 cells demonstrated significant cellular growth reduction upon metabolic rewiring by exogenous administration of potassium lactate. Clinical studies have shown the crucial roles of IFN-γ for successful cancer immunotherapies involving checkpoint inhibitors and chimeric antigen receptor T cells. The positive implications of this study on the metabolic modulation of IFN-γ activation on heterogeneous tumour cells are discussed.

## Introduction

IFN-γ is the sole member of type II interferons primarily produced by cells of the immune system, such as T cells and natural killer (NK) cells. IFN-γ is essential in tissue homeostasis, inflammatory responses, and tumour immunosurveillance [1]. IFN-γ is involved in many inflammatory disease progression and pathogenesis. A lack of IFN-γ receptor 1 causes severe mycobacterial infections and viral infections caused by herpes viruses, parainfluenza virus type 3, and respiratory syncytial virus, with a significant risk of early mortality [2]. Immunocompromised children with IFN-γ signalling deficiency are highly susceptible to lethal disseminated bacillus Calmette-Guerin: a condition known as BCGosis and local recurrent nontuberculous mycobacterial infection [3,4]. Other infectious diseases like leishmaniasis, AIDS, malaria, influenza, COVID-19, etc., are also alleviated through the IFN-γ-signalling processes [5]. However, IFN-γ-hyperactivation negatively impacts several inflammatory and autoimmune diseases: multiple sclerosis, rheumatoid arthritis, encephalomyelitis, Sjogren’s syndrome, and inflammatory bowel disease, among others [6–10].

The canonical IFN-γ signalling triggers the JAK-STAT pathway involving JAK2, MEK1/2, Erk1/Erk2, and STAT1α as vital players [11]. STAT1α begins transcribing a set of primary response genes, followed by IRF1-mediated secondary response gene expression [12]. Several transcription factors from the IFN-γ response, like IRF1 and NF-κB, bind to the *Nos2* promoter and synergistically regulate *Nos2* expression [11,13]. NOS2, an isoform of the NOS enzymes, biosynthesizes Nitric Oxide (NO) from L-arginine and requires several co-factors: reduced nicotinamide-adenine-dinucleotide phosphate, flavin adenine dinucleotide, flavin mononucleotide, and (6R-)5,6,7,8-tetrahydrobiopterin [14]. The cellular oxidative state greatly influences the catalytic efficiency of the NOS enzymes. The expression of NOS2 catalysing the production of NO is one of the key markers in IFN-γ signalling in different cell types, including macrophages and tumours [1,15,16]. NO, a short-lived bioactive gaseous molecule, plays pleiotropic physiological roles in normal cells and pathophysiological roles in cancer. NO is a component of the IFN-γ-signalling response in macrophages and tumours [15–17]. IFN-γ signalling induces the transcription of *Nos2* that upregulates the intracellular NOS2 amounts. NOS2 is responsible for the heightened NO production in IFN-γ-activated cancer cells. IFN-γ-modulated genes and responses can be divided into two groups: oxidative and nitrosative stress dependent and oxidative and nitrosative stress independent [15,16].

The amounts of IFN-γ in the tumour microenvironment, along with the cellular, microenvironmental, and molecular contexts, play major role in the anti-tumour response. Tumours regress under high amounts of IFN-γ through apoptosis and ferroptosis [18] and recombinant IFN-γ therapy against adult T-cell leukaemia is approved in Japan [19]. Clinical investigations showed that intact IFN-γ signalling is essential for successful immunotherapy against cancer. A genome-wide CRISPR knockout screen in glioblastoma identified that components of the IFN-γ signalling cascade are crucial for the CAR-T cell-mediated killing of tumours [20]. Also, clinical responses to immune checkpoint blockade therapy against tumours like melanoma require functional IFN-γ signalling components for tumour regression [21,22]. Therefore, understanding the tumoral response to IFN-γ-signalling is important for understanding host response during cancer immunotherapy.

The progression and functions of IFN-γ signalling are intricately associated with metabolic reprogramming. The production of IFN-γ is tightly regulated by T-cell metabolism, where upregulation of glycolysis favors IFN-γ transcription and translation. GAPDH binds to the 3’-UTR of IFN-γ mRNA to negatively regulate IFN-γ production in naive T-cells [23]. IFN-γ production by NK cells upon IL15 activation also depends on elevated glycolysis [24]. IFN-γ-mediated classical activation of macrophages toward an inflammatory state requires elevated glycolytic flux and limited reliance on mitochondrial oxidative metabolism [25]. IFN-γ-mediated enhanced aerobic glycolysis is essential for clearing Mycobacterial infection in mice [26]. These studies led us to investigate the metabolic changes in heterogeneous tumours during IFN-γ activation. Previous studies from our group demonstrated that IFN-γ-activated mouse tumour cells, such as H6 hepatoma and L929 fibrosarcoma, undergo cell cycle arrest and apoptosis [15,16]. In this study, we utilized the *in vitro* model of the IFN-γ-activation of tumour cells to address whether the tumour growth arrest relied on metabolic alterations and dissected the underlying mechanism to identify the key regulatory factors. Finally, we incorporated these key regulatory factors into tumours that do not undergo growth arrest upon IFN-γ-activation. This novel approach led to the growth arrest of the tumours with IFN-γ-activation. It has raised the potential of targeting metabolic processes downstream to IFN-γ-activation for an enhanced and effective immunotherapy response against tumours.

## Materials and Methods

### Instruments and reagents

The list of instruments and reagents along with country of origin and catalogue numbers are listed in supplementary Table 1.

### Cell culture

H6 (hepatoma), Renca (renal adenocarcinoma), RAW 264.7 (monocyte/macrophage), CT26 colon carcinoma, and B16F10 melanoma cell lines were maintained in Dulbecco’s modified minimal essential medium (DMEM), with 10% FBS [27]. The media was supplemented with 5 μM β-mercaptoethanol, 100 μg/ml penicillin, 250 μg/ml streptomycin, 50 μg/ml gentamycin, and 2 mM glutamine. The cells were maintained at 37°C in a humidified incubator with 5% CO_2_ and 95% humidity. More detailed features of the cell lines used in this study are tabulated in Supplementary Table 2.

### Nitrite measurement

NO production was estimated by measuring the levels of accumulated nitrite, the metabolic product of NO metabolism, in the medium using Griess reagent [28]. The reagent was prepared as a mixture of 1% (w/v) Sulfanilamide and 0.1% (w/v) N-(1-Naphthyl) ethylenediamine dihydrochloride dissolved in 2.5% ortho-phosphoric acid. The reaction mixture for the assay consisted of 25 μL of the cell-free supernatant, 25 μl of 10% DMEM, and 100 μl of Griess reagent that were mixed well. The absorbance was measured at 550 nm using the Tecan microtiter plate reader. The amount of nitrite in the supernatants was calculated from a standard curve of sodium nitrite (1.22 – 625 μM).

### Trypan blue dye exclusion assay

Trypan blue dye exclusion assay was performed to assay for percentage change in cell number [16]. Briefly, logarithmically growing cells were seeded at an initial density of 10^4^ cells per well in a 96-well plate. The cells were treated with IFN-γ and harvested by adding 0.5% Trypsin-EDTA (HiMedia, India) per well. The cell suspension was mixed with an equal volume of 0.4% Trypan blue, and the number of live cells was counted in a hemacytometer.

### Seahorse XF analyses

Glycolysis Stress Test and Mitostress test were done to measure the ECAR and OCR from H6 cells. ∼0.8x10^4^ H6 cells were seeded in an 8-well plate in each well, allowed to adhere for 8 h, and treated with IFN-γ for 24 h before being subjected to XF Analyzer (Seahorse Biosciences). The experiments were performed according to the manufacturer’s protocol, using the reagents (10 mM D-glucose, 1 mM oligomycin, 50 mM 2-Deoxy D-glucose) for the glycolysis stress test and (1.5 μM oligomycin, 1 μM FCCP, 0.5 μM Rotenone, and Antimycin A) for the mitostress test. The manufacturer supplied the XF media, which was supplemented with 2 mM L-glutamine.

### Colorimetric assay for glucose and lactate

Glucose uptake and accumulated lactate were estimated from the cell-free supernatant using a colorimetric glucose estimation kit and lactate assay kit by slightly modifying the manufacturer’s protocol to adapt for a microtiter plate reader-based readout. Briefly, for glucose uptake assay, 5 µL of cell-free supernatant, 15 µL of distilled water, and 200 µL of the reaction mix were mixed and incubated for 15 minutes at room temperature. Subsequently, absorbance was measured at 340 nm using a Tecan microtiter plate reader. The cell-free supernatant was diluted in the assay buffer in a 1:1000 dilution for lactate assay. 50 μL of the reaction mixture was added to 50 μL of the diluted supernatant and incubated for 30 minutes at 37°C. The absorbance was recorded at 540 nm at an interval of 10 minutes for lactate assay using a Tecan microtiter plate reader.

### RNA isolation and quantitative real-time PCR

RNA isolation from H6 cells was performed as previously described [29]. H6 cells were seeded in 6 well plates at a density of 3 × 10^5^ cells per well. The cells were treated with IFN-γ (10 U/mL) and harvested kinetically. The cell lysates were prepared using TRI reagent. Phenol-chloroform extraction of RNA was performed. The mRNA from 1-3 μg total RNA was reverse transcribed to cDNA using a first-strand cDNA synthesis kit. qRT-PCR was performed with the following program: initial denaturation (95°C for 10 minutes), denaturation (95°C for 10 seconds), annealing (57°C for 30 seconds), and extension (70°C for 20 seconds). The denaturation to extension steps was repeated for 39 cycles. The C_t_ values of gene expression data were transformed to 2^-ΔΔC^_t_ and represented in the figures. The primer sequences are listed in supplementary Table 3.

### Flow cytometry (2-NBDG, ROS, TMRE, MHC class 1)

Glucose uptake and mitochondrial membrane potential were studied using 2-NBDG [30] and TMRE [31] dyes. The intracellular ROS estimation and surface staining of MHC class 1 were performed as previously described [16]. Briefly, H6 cells were cultured in 6-well plates at a density of 3 × 10^5^ cells per well and treated with IFN-γ. Cells were incubated with 100 µM 2-NBDG dye in PBS at 37°C water bath for 30 mins in the dark to estimate glucose uptake. Cells were incubated with 150 nM TMRE dye in serum-free DMEM in the dark at room temperature for 10 mins to assess mitochondrial membrane potential. The intracellular ROS was estimated by incubating the cells with 10 µM DCFDA dye at 37°C water bath for 30 mins in the dark. Cell pellets were resuspended in 200 μL of PBS in all three cases. Cell surface staining for MHC class 1 was performed by incubating the cells in the blocking buffer {5% FBS and 0.1% (w/v) sodium azide in PBS} for 30 min. The cells were stained with the PE-conjugated anti-MHC Class 1 antibody at the dilution 1:300 for 30 mins with intermittent tapping. The cells were fixed with 4% paraformaldehyde. Data acquisition was performed in the BD FACSVerse™ flow cytometer and analyzed using BD FlowJo™ (BD Biosciences US).

### Western Blot

Western blot was performed as described [32]. 0.5-0.6 million H6 cells were collected, washed in PBS, and lysed in RIPA lysis buffer containing protease and phosphatase inhibitors. The lysates were centrifuged at 14000g for 30 minutes to remove debris, and the supernatant was collected. Protein concentration was determined using the BCA assay. Next, the proteins were separated by SDS-PAGE using a polyacrylamide gel, followed by electrotransfer onto a PVDF membrane in a semi-dry transfer apparatus. The membrane was then blocked with a blocking solution (e.g., 5% non-fat skim milk in TBST) for 30 minutes on a rocker to prevent non-specific binding. After blocking, the membrane was incubated overnight with a primary antibody specific to the target protein (1:5,000 dilution for HIF-1*α* or β-Actin antibody, diluted in blocking buffer) at 4°C. Subsequently, the membrane was washed with TBST for 30 minutes at room temperature to remove the unbound primary antibody and then incubated with the secondary antibody conjugated to HRP (1:10,000 dilution in TBST) for 2 hours at room temperature on a rocker. After three washing rounds, the target protein bands were visualized using a chemiluminescent HRP substrate in chemidoc instrument. The band intensities were quantified using Multigauge V3.0 software, and the results were analyzed to determine protein expression levels.

### Statistical analyses

All graphical representations and statistical analyses were performed on GraphPad Prism software version 8.0.2. All data are represented as mean ± standard deviation of the mean (SD) unless stated otherwise. The data distribution was tested using QQ-plot to examine the skewness. Statistical analyses were performed using ordinary One-way ANOVA with Sidak’s multiple comparisons tests and two-way ANOVA with Tukey’s multiple comparisons tests. (ns), (*), (**), (***), (****) indicate non-significant difference and the statistical differences of p < 0.05, p < 0.01, p < 0.001, and p<0.0001, respectively, between the comparable indicated, unless stated otherwise.

## Results

### IFN-γ activation increases NO and glycolysis-mediated extracellular acidification with growth reduction in H6 cells

Previous research from our group demonstrated that H6 cells undergo cell cycle arrest and apoptosis upon activation by IFN-γ, which is dependent on intracellular NO and ROS [15,16]. In this study, we attempted to test whether the cytostatic fate of tumours upon IFN-γ-activation is associated with metabolic reprogramming. The cellular respiratory and metabolic alterations often lead to the release of gaseous byproducts and metabolites that can influence the extracellular pH [33,34]. Initially, we used the H6 cells as a model to study IFN-γ activation [15,16] and examined the pH of the cell culture medium. The cell culture medium contains phenol red as a pH indicator. After 24 hours of IFN-γ-activation, the cell-free supernatant of H6 cells was visibly more orange compared to the untreated control (Fig.S1B). The phenomenon did not cause any noticeable cytomorphological change in the H6 cells (Fig.S1A). The pH measurement revealed that the mean pH of cell-free supernatant from IFN-γ-treatment was 6.64±0.03, less than the untreated control, which was 6.84±0.02 (Fig.S1C). The pH scale is logarithmic, meaning that a change of 0.2 corresponds to a twofold increase in acidity, which may be physiologically significant.

One possibility was that the increased cell growth may enhance extracellular acidification upon IFN-γ-activation. The nitrite and lactate production significantly increased post-24 hours of IFN-γ-activation. However, the total cell number reduction was not significant due to variations in data points despite an overall trend (Fig.S1D). Therefore, the data was expressed as the percent cell number change and the statistically significant cell number reduction was obtained after 24 hours of activation [15,16]. The *in vitro* acidification of IFN-γ-activated H6 cells was also accompanied by a kinetic increase in nitrite production. The nitrite amounts were normalized to the percent cell number values for a fair analysis. The level of nitrite significantly increased after 12 and 24 hours of IFN-γ-activation (Fig.1A).

**Fig.1.**
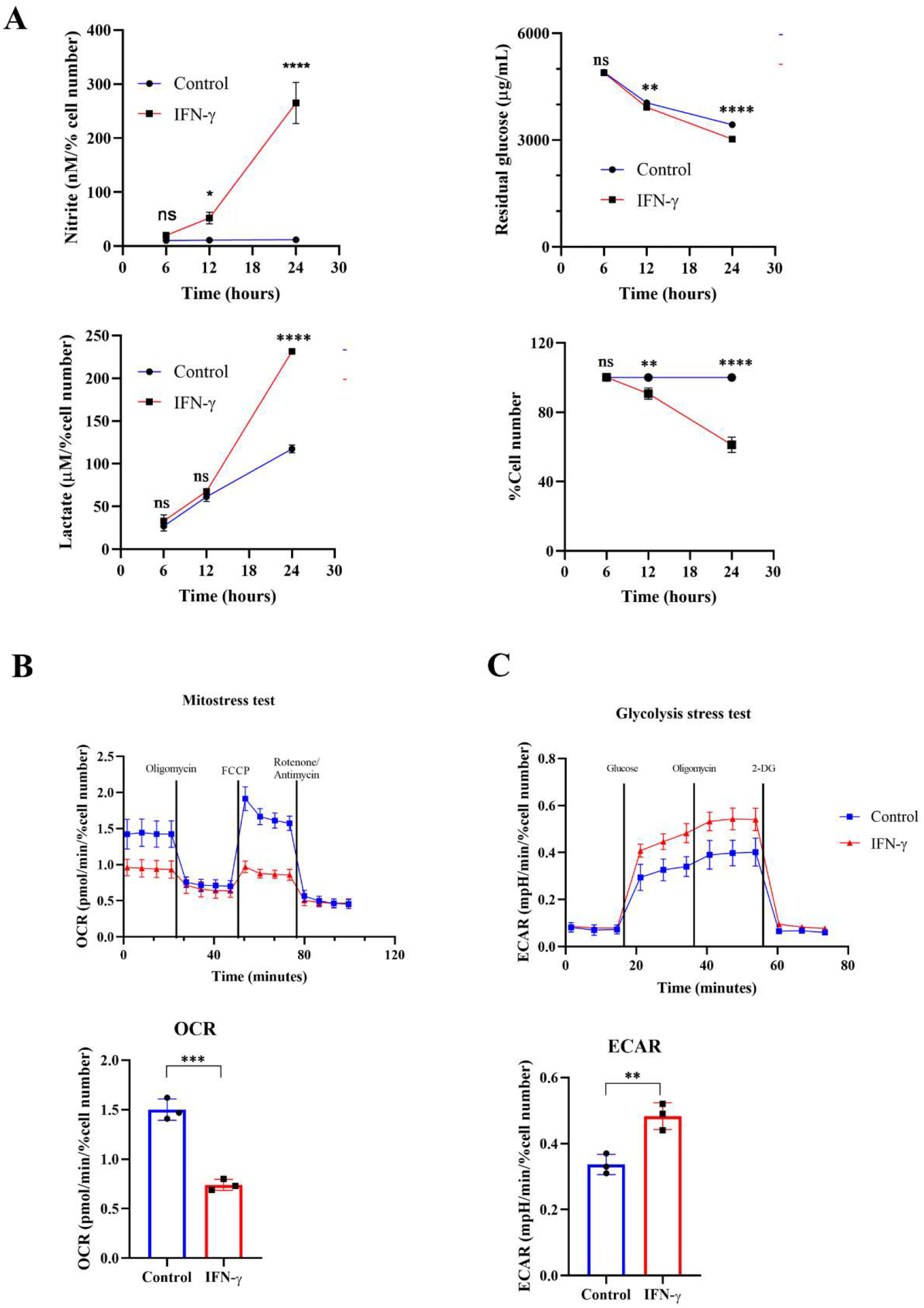
IFN-γ-activation of H6 cells increases glycolytic flux and compromises mitochondrial function. H6 hepatoma cells were seeded in a 6-well plate and treated with 10 U/mL IFN-γ for the indicated time points. The cell numbers were counted, and the cell-free supernatant was collected and assayed for nitrite, residual glucose, and lactate. The nitrite and lactate levels were normalized to the percent cell number (A). The Seahorse XF analysis of the H6 cells was performed post-IFN-γ-activation for 24 hours. The pharmacological modulators were sequentially injected as indicated. The mitostress test with the OCR from the 4th measurement (B) and the glycolysis stress test with the ECAR from the 3rd measurement (C) are shown. The statistical analyses were performed using two-way ANOVA with Tukey’s multiple comparisons tests (A) and unpaired t-tests (A-C). (ns), (*), (**), (***), (****) indicate non-significant difference and the statistical differences of p < 0.05, p < 0.01, p < 0.001, and p<0.0001 between the comparable indicated. Each data point is representative of an independent experiment. Data are represented as mean ± SD of 3-5 independent experiments.

It was important to address the source of extracellular acidification during IFN-γ-activation of H6 cells. We used multiple approaches to address this aspect: the TMRE-based flow cytometric assessment tested mitochondrial membrane potential and the seahorse mitostress test assessed mitochondrial oxygen consumption rate (OCR). IFN-γ-activation of H6 cells significantly reduced mitochondrial membrane potential and OCR. The reduced OCR in IFN-γ-activated H6 cells was significantly less responsive to pharmacological inhibitors of mitochondrial function, i.e., oligomycin, FCCP, rotenone, and antimycin (Fig.1B). Therefore, these assessments indicated that the compromised mitochondrial function might not increase CO_2_ release upon IFN-γ-activation.

We further tested whether the glycolytic mean of metabolic acidosis contributes to the IFN-γ-induced enhanced extracellular acidification. The glycolysis stress test revealed that glucose-starved IFN-γ-activated H6 cells display a significantly heightened extracellular acidification rate (ECAR) upon glucose injection. The ECAR enhancement did not alter upon F_o_-F_1_ inhibition by oligomycin, proving no involvement of mitochondrial ATP production. The inhibition of hexokinase dampened ECAR, ultimately levelling the untreated H6 cells (Fig.1C). In conclusion, the IFN-γ-mediated heightening of ECAR remarkably depended on glycolysis. The H6 cells consumed glucose and produced lactate in significantly higher amounts after 24 hours of IFN-γ-activation (Fig.1A). Therefore, IFN-γ-activation rewired the cellular metabolism to augment the glycolytic production of lactate and skewed mitochondrial functions.

### IFN-γ activation boosts NO and ROS formation, enhancing glycolytic flux and upregulating lactate production from H6 cells

Next, we asked whether NOS2-induced NO plays any regulatory role in enhancing the IFN-γ-mediated glycolytic flux [15]. The pharmacological inhibitors of NOS function, i.e., *N*^ω^-Methyl-L-arginine (LNMA; inhibits all NOS isoforms) or 1400W (specifically inhibits NOS2), were added in the presence of IFN-γ [15,16,35]. Both the NOS inhibitors significantly reduced the IFN-γ-induced NO and ROS production and rescued cell growth reduction (Fig.2A, D; S2A). The efficiency of glucose uptake was tested using 2-NBDG-based flow cytometry and measuring the residual glucose amounts in the cell-free supernatant. NOS inhibition significantly reduced the glucose uptake and lactate production of IFN-γ-activated H6 cells (Fig.2B-C, S2B). Therefore, IFN-γ-induced NO production dominantly increases the glycolytic flux in H6 cells.

**Fig.2.**
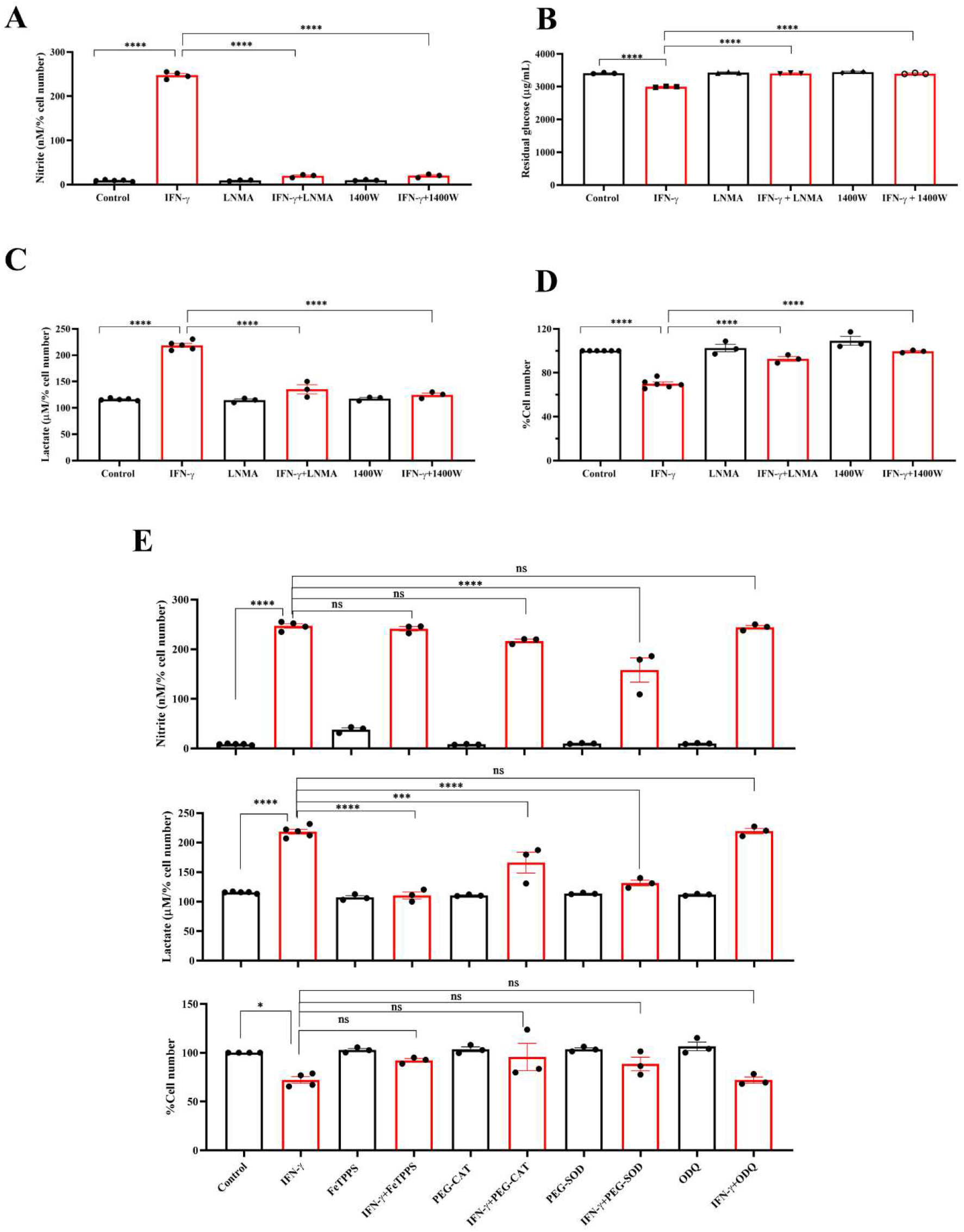
IFN-γ-activated augmentation of glycolytic flux is NO and ROS-dependent in H6 cells. H6 cells were treated with 10 U/ml IFN-γ and incubated alone or with pharmacological inhibitors of NOS enzymes (LNMA at 200 uM and 1400W at 6 uM) for 24 hours to inhibit NO biosynthesis. The nitrite levels were measured from the cell-free supernatants and normalized to the percent cell number (A). The residual glucose concentrations in the cell-free supernatant were measured after 24 hours of IFN-γ-activation of the H6 cells (B). The lactate concentrations were measured in the same cell-free supernatant and normalized to percent cell number (C). The cell numbers were counted and represented as the percent cell number (D). The H6 cells were treated with 10 U/ml IFN-γ and incubated alone or with the peroxynitrite quencher (FeTPPS at 50 uM), inhibitors of ROS production (PEG–CAT at 100 U/ml and PEG–SOD at 50 U/ml) and soluble guanylate cyclase inhibitor ODQ at 20 uM for 24 hours. The cell numbers were counted and represented as the percent cell number. The nitrite and lactate concentrations were measured in the cell-free supernatant and normalized to percent cell number. (E). The statistical analyses were performed using ordinary one-way ANOVA with Sidak’s multiple comparisons tests. (ns), (*), (**), (***), (****) indicate non-significant difference and the statistical differences of p < 0.05, p < 0.01, p < 0.001, and p<0.0001 between the comparable indicated. Each data point is representative of the independent experiment. Data are represented as mean ± SD of 3-5 independent experiments.

Subsequently, we asked whether lactate production can be enhanced by further elevation of intracellular NO. SNAP, a NO-donor compound, was added to the H6 cells in the presence of IFN-γ. The exogenous NO-donation approach did not elevate the IFN-γ-induced production of nitrite.

The level of lactate and percent cell number remained unaffected (Fig.S3A). Therefore, we opted for an alternative strategy involving the concept of ‘arginine paradox’. In this phenomenon, increased arginine availability leads to increased NO production despite a theoretical intracellular saturation of L-arginine for the NOS enzyme activity [36]. H6 cells were treated with L-arginine in varying concentrations in the presence of IFN-γ for 24 hours. This approach successfully elevated IFN-γ-induced NO production in a concentration-dependent manner. However, the IFN-γ-induced increased lactate production and reduced cell growth remained unaffected despite the significant increase in NO production (Fig.S3B, C). These results suggest that the NO-mediated elevation of lactate production occurs at maximum saturation upon the IFN-γ-activation of H6 cells.

One of the mechanisms of the effects of NO is exerted by activating intracellular soluble guanylate cyclase (sGC) and subsequent upregulation of cyclic GMP (cGMP) [37]. We asked whether the cGMP upregulation is responsible for enhancing lactate release. ODQ, a pharmacological inhibitor of sGC, did not affect the IFN-γ-induced nitrite and lactate production and reduction in percent cell number (Fig.2E). Therefore, cGMP-activation was not involved in the IFN-γ-mediated enhancement of lactate production. Next, we asked whether NO-induced ROS levels affect the lactate release in the IFN-γ-activated H6 cells [16]. Peroxynitrite, hydrogen peroxide, and superoxide radical levels were quenched using FeTPPS, pegylated catalase (PEG-CAT), and pegylated superoxide dismutase (PEG-SOD), respectively. PEG-SOD partially but significantly reduced NO production, indicating that the oxidative environment created by superoxide radicals promoted NO biosynthesis. The IFN-γ-mediated cell growth reduction was partially but non-significantly rescued in the presence of all three RNS and ROS quenchers. All three RNS and ROS quenchers significantly reduced the IFN-γ-mediated lactate production (Fig.2E). In conclusion, IFN-γ-mediated NO and subsequent ROS production are responsible for IFN-γ-mediated increased lactate production in H6 cells.

### IFN-γ-activated glycolytic flux augmentation promotes NO formation and reduces cellular growth of H6 cells

The IFN-γ-mediated enhancement of lactate release raised the question of why the cells undergo metabolic reprogramming toward the induction of glycolysis and what the cells gain from the glycolytic dependency. The question was addressed using pharmacological inhibition of several nodes of the glycolytic pathway: 2-deoxyglucose (2-DG, an inhibitor of hexokinase), GSK2837808A (an inhibitor of lactate dehydrogenase A or LDHA), and AR-C155858 (an inhibitor of monocarboxylate transporter 1-4 or MCT1-4). 2-DG, GSK2837808A, and AR-C155858 block the first step of glycolysis, lactate production, and lactate shuttling in and out of the cells, respectively. All the inhibitors significantly reduced glucose uptake and lactate production and release by the IFN-γ-activated H6 cells (Fig. 3C, E; S4B). However, none of the inhibitors affected the IFN-γ-mediated induction of MHC class 1 surface expression and reduction in mitochondrial membrane potential (Fig. 3D; S4A, C). All the inhibitors partially but significantly reduced the IFN-γ-induced NO production (Fig. 3A) and 2-DG significantly reduced the IFN-γ-induced ROS production (Fig. 3B). Importantly, all the inhibitors partially but significantly rescued the IFN-γ-mediated percent cell number reduction (Fig. 3F). In conclusion, the IFN-γ-activated H6 cells produce NO and ROS to enhance glycolysis. Most likely, the cells harness the utilities of IFN-γ-induced glycolytic flux augmentation to promote NO production further and reduce cellular growth.

**Fig.3.**
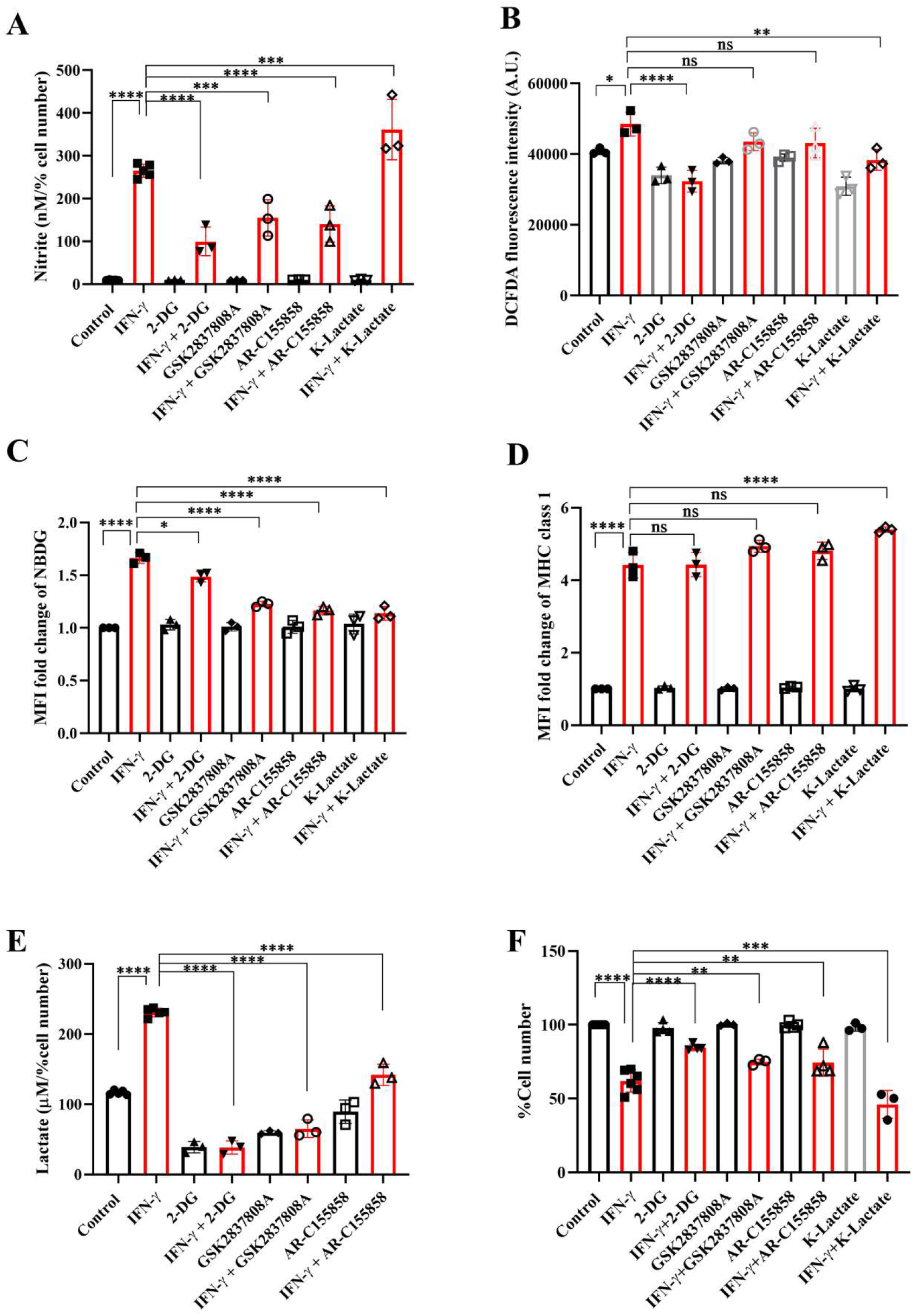
IFN-γ-mediated enhanced glycolysis fuels IFN-γ signalling responses by increasing NO and decreasing cell number. H6 cells were treated with 10 U/ml IFN-γ and incubated alone or with pharmacological modulators of glycolysis (2-DG, a hexokinase inhibitor at 2.5 mM; GSK2837808A, a lactate dehydrogenase inhibitor at 125 nM; AR-C155858, a monocarboxylate transporter inhibitor at 6 nM and K-lactate as a metabolic reprogramming agent at 10 mM). The cell-free supernatants were collected after 24 hours, and the level of nitrite and lactate were estimated (A, E). The cell numbers were counted to derive the percent cell number (F). The levels of nitrite and lactate were normalized to the percent cell number. The intracellular ROS was measured using a Tecan microplate reader from 0.1 million cells/well upon DCFDA staining for 30 minutes (B). The IFN-γ-activated H6 cells were assayed for glucose uptake and surface expression of MHC class 1 using flow cytometry in the presence of the glycolysis modulators. The cells were incubated with 100 μM 2-NBDG for 30 minutes, and flow cytometric analysis of glucose uptake was performed (C). The cells were stained with an antibody to MHC class 1, and flow cytometric analysis of the surface expression of MHC Class 1 was performed (D). The statistical analyses were performed using ordinary one-way ANOVA with Sidak’s multiple comparisons tests. (ns), (*), (**), (***), (****) indicate non-significant difference and the statistical differences of p < 0.05, p < 0.01, p < 0.001, and p<0.0001 between the comparable indicated. Each data point is representative of the independent experiment. Data are represented as mean ± SD of 3 independent experiments.

To address the role of lactate in the IFN-γ signalling responses, potassium lactate was added along with IFN-γ to the H6 cells. Potassium lactate did not reduce the pH of the cell culture medium and was preferred over lactic acid to avoid the effects of pH reduction [38]. Potassium lactate, upon IFN-γ-activation, completely rescued the extracellular acidification and pH reduction, from 6.65±0.04 in IFN-γ to 6.97±0.05 in the combination of IFN-γ and potassium lactate. Potassium lactate significantly reduced the IFN-γ-mediated enhancement of intracellular ROS and glucose uptake (Fig. 3B, C; S4B). These indicate that potassium lactate reduced the glucose dependency of the IFN-γ-activated cells and most likely, served as an alternative carbon source to rewire the metabolism. Notably, the rewired metabolism significantly augmented the IFN-γ-induced NO production and cell growth reduction (Fig. 3A, F). These observations indicate that the IFN-γ-activated H6 cells may utilize the heightened production of lactate to promote IFN-γ signalling in an autocrine manner.

### IFN-γ promotes HIF-1***α*** stabilization and transcriptional upregulation of HIF-1***α*** target glycolytic genes in H6 cells

Previous reports suggested that pharmacological donation of NO and elevation of intracellular ROS can activate the hypoxia signalling process by inhibiting the prolyl hydroxylase (PHD) enzyme. Also, HIF-1*α* is a well-known regulator of glycolytic flux in tumour cells [39]. Therefore, we tested whether the IFN-γ-activation of H6 cells stabilizes HIF-1*α* and upregulates glycolytic gene expression. RT-qPCR-based gene expression analysis revealed that IFN-γ-activation significantly upregulated several well-known members of IFN-γ-activated gene expression. The transcription of *Irf1* and *Nos2* was significantly upregulated after 6 and 12 hours of activation. *Cd274* was significantly upregulated after 6 hours of IFN-γ-activation. Along with the well-known gene expression profile, IFN-γ-activation also significantly upregulated *Hif1a* transcription after 6 and 12 hours (Fig.4A). Immunoblot analysis of HIF-1*α* revealed that the HIF-1*α* levels were significantly increased after 12 hours of IFN-γ-activation in H6 cells (Fig. 4B), indicating that the upregulation of intracellular HIF-1*α* is regulated at both mRNA and protein levels.

**Fig.4.**
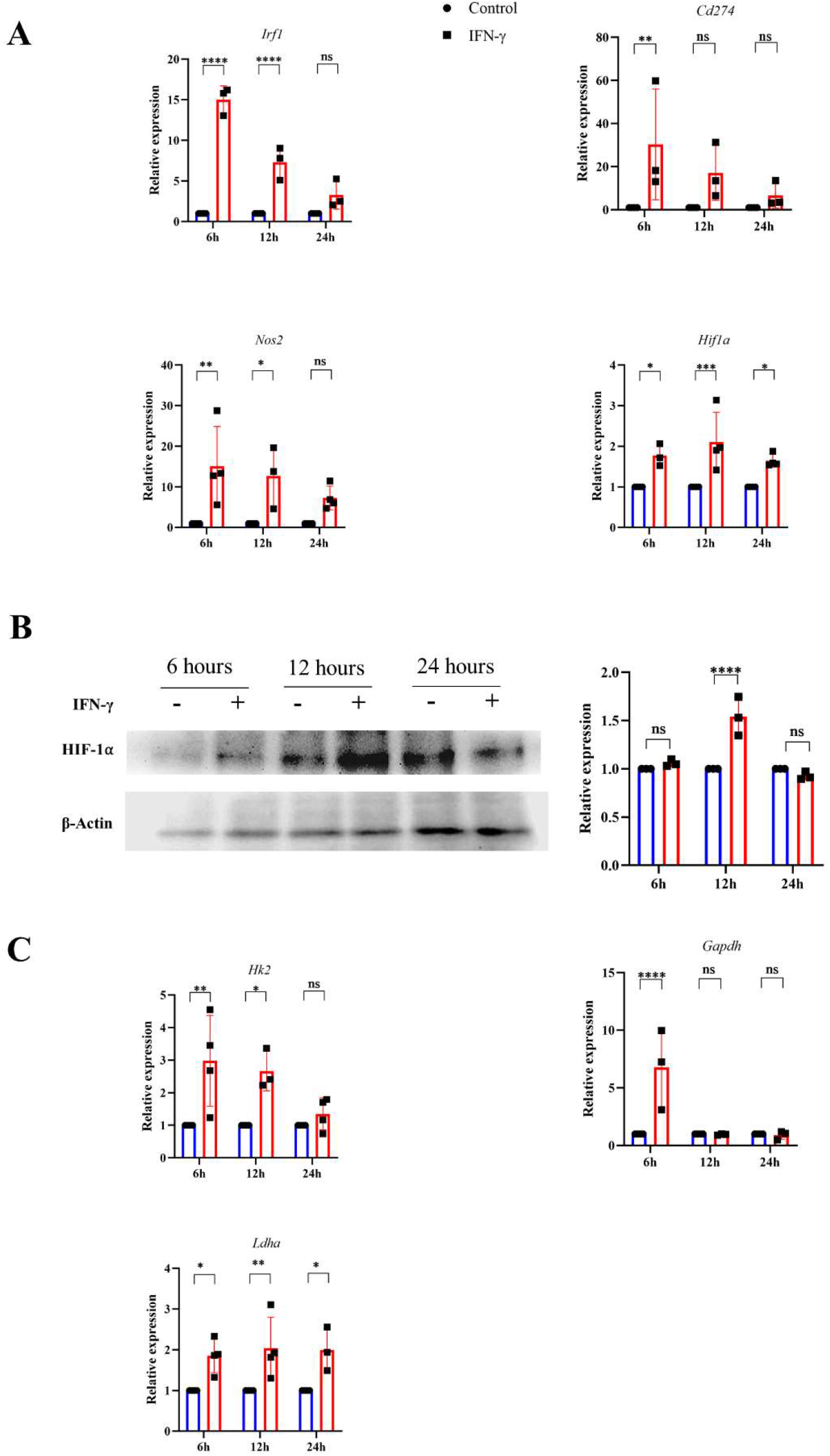
IFN-γ-activation stabilizes HIF-1α and induces several HIF-1α-responsive glycolytic genes. H6 cells were treated with 10 U/ml IFN-γ in a 6-well plate for the indicated time points, and total RNA was extracted. RT-qPCR was performed to quantify the relative expression of *Irf1*, *Nos2*, *Cd274*, and *Hif1a* (A). The cells were lysed at the indicated time, and immunoblot was performed to quantify the HIF-1*α* intracellular protein amounts. β-Actin was used as the loading control (B). The relative mRNA expression of several HIF-1*α* target genes, such as *Hk2*, *Gapdh*, and *Ldha*, was also quantified using RT-qPCR (C). The statistical analyses were performed using ordinary two-way ANOVA with Tukey’s multiple comparisons tests. (ns), (*), (**), (***), (****) indicate non-significant difference and the statistical differences of p < 0.05, p < 0.01, p < 0.001, and p<0.0001 between the comparable indicated. Each data point is representative of the independent experiment. Data are represented as mean ± SD of 3-4 independent experiments.

A list of HIF-1*α*-induced differentially upregulated genes (log_2_ fold change ≥ 1.5) was obtained from the NCBI GEO dataset (GSE98060) [40], analysed with GEO2R, and imported into the String tool of EMBL-EBI. A protein-protein interaction (PPI) network was constructed using the String tool to identify the functional enrichment of the topmost interconnected cluster and Cytoscape software analysed the network. The network had 354 nodes, 48 edges, an average node degree of 0.271, with an average local clustering coefficient of 0.0998. KEGG pathway analysis showed that the glycolysis pathway is the second most enriched pathway after the HIF-1*α* signalling pathway (Supplementary table 4). The K-means method and MCODE-based computation of network clustering identified that the topmost interconnected regulatory network of HIF-1*α* majorly consists of several glycolytic genes (Fig.S5A, B). Based on these data, we asked whether the IFN-γ-induced glycolytic flux enhancement is associated with the transcriptional upregulation of HIF-1*α*-target glycolytic genes: *Hk2*, *Gapdh*, and *Ldha*. *Hk2*, *Gapdh*, and *Ldha* encode hexokinase 2, glyceraldehyde-3-phosphate dehydrogenase, and lactate dehydrogenase a, respectively. All three glycolytic genes were significantly upregulated after 6 hours of IFN-γ-activation. *Hk2* and *Ldha* were significantly upregulated after 12 hours of IFN-γ-activation (Fig. 4C). HIF-1*α* is known to upregulate *Vegfa* to promote tumour angiogenesis. However, the RT-qPCR-based analysis revealed that *Vegfa* was not differentially regulated in IFN-γ-activated H6 cells (Fig. S6C). These indicate that the IFN-γ-induced HIF-1*α*-mediated gene expression does not mimic the canonical hypoxia-induced HIF-1*α* gene expression signature but involves specific ones, especially glycolytic genes. Lactate dehydrogenase a (LDHA) and Lactate dehydrogenase b (LDHB) perform opposite functions. LDHA promotes pyruvate oxidation to lactate, whereas LDHB promotes lactate reduction to pyruvate [41]. *Ldhb* was also not differentially regulated upon IFN-γ-activation, whereas *Ldha* was significantly upregulated 24 hours post-activation by IFN-γ (Fig. 4C; S6C). These observations indicate that the glycolytic flux upregulation is accompanied by *Hif1a* expression, HIF-1*α* stabilization, and HIF-1*α*-targeted glycolytic gene expression.

### The IFN-γ-induced function of HIF-1***α*** promotes NO and ROS production, enhances glycolytic flux, and lowers cellular growth

The functional roles of HIF-1*α* in the IFN-γ-induced glycolytic flux was addressed using pharmacological approaches: chetomin (an inhibitor of HIF-1*α*) and DMOG (a pharmacological HIF-1*α* stabilizer) were used in IFN-γ signalling. The HIF-1*α* stabilizer DMOG did not affect the IFN-γ-induced NO and ROS production, mitochondrial membrane potential, MHC class 1 surface expression, glucose uptake, and cell growth reduction (Fig.5A-D, F; Fig. S7). However, DMOG significantly increased IFN-γ-induced lactate release, indicating that the IFN-γ-activated H6 cells contain more room for glycolytic flux enhancement (Fig. 5E). The HIF-1*α* inhibitor chetomin reduced IFN-γ-induced NO and ROS production and glycolytic flux and rescued cell growth reduction significantly (Fig.5A-E; S7C-D). Hence, these observations demonstrate that HIF-1*α* partially regulates NO and ROS production, whereas it strongly regulates the IFN-γ-induced glycolytic flux enhancement. More importantly, HIF-1*α* significantly rescued IFN-γ-induced cell growth reduction (Fig. 5F). Therefore, HIF-1*α* stabilization substantially enhances glycolytic flux downstream of NO, thereby reducing cell growth.

**Fig.5.**
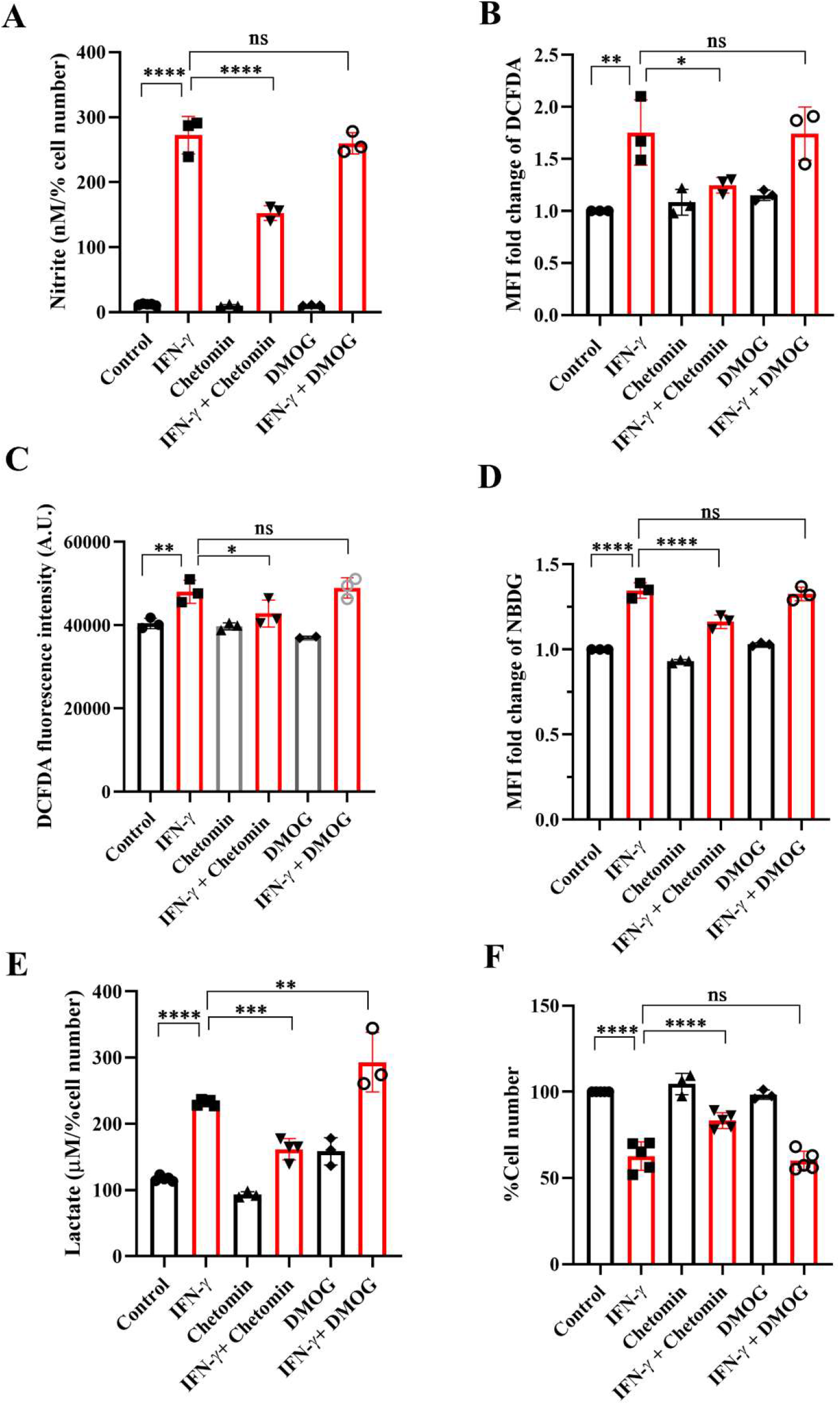
IFN-γ-activated augmentation of glycolytic flux is dependent on HIF-1α in H6 cells. H6 cells were treated with 10 U/ml IFN-γ for 24 hours. The IFN-γ-activated cells were incubated alone or with pharmacological modulators of HIF-1*α* function (Chetomin, an inhibitor of HIF-1*α* at 12.5 nM, and DMOG, a stabilizer of HIF-1*α* at 0.5 uM). The cell-free supernatant was collected and the levels of nitrite and lactate were estimated in the cell-free supernatant. The nitrite and lactate levels are normalized to percent cell number (A, E). The IFN-γ-activated H6 cells were assayed for intracellular ROS by incubating the cells with 10 μM DCFDA for 30 minutes. The analyses of intracellular ROS were performed using flow cytometry (B) and Tecan microplate reader-based estimation (C). The cells were incubated with 100 μM 2-NBDG for 30 minutes, and flow cytometric analysis of glucose uptake was performed (D). The cell number was counted to derive the percent cell number (F). The statistical analyses were performed using ordinary one-way ANOVA with Sidak’s multiple comparisons tests. (ns), (*), (**), (***), (****) indicate non-significant difference and the statistical differences of p < 0.05, p < 0.01, p < 0.001, and p<0.0001 between the comparable indicated. Each data point is representative of the independent experiment. Data are represented as mean ± SD of 3-5 independent experiments.

### Lactate production induced by IFN-γ through NO is observed in other NO-producing cells

The NO-dependent regulation of glycolytic flux enhancement upon IFN-γ-activation of H6 cells led us to ask whether the process is also observed in other tumour cell lines. Raw 264.7 (monocyte/macrophage) and Renca (renal adenocarcinoma) tumour cell lines produced NO and reduced cell number post IFN-γ activation, phenocopying the response of the H6 cell line. IFN-γ-induced Raw 264.7 and Renca cells also produced heightened amounts of lactate.

### The addition of exogenous potassium lactate in the presence of IFN-γ decreases cellular growth in non-NO-producing cells

Two non-NO-producing cell lines CT26 (colon carcinoma) and B16F10 (melanoma) did not increase NO and reduce cell growth upon IFN-γ-activation. None of the non-NO-producing cell lines increased lactate; however, B16F10 showed slight but significant reduction in lactate production (Fig.6). Therefore, we asked whether the reconstitution of the IFN-γ-inducible missing components, such as NO, HIF-1α, and lactate, can sensitize the resistant non-NO-producing cell lines into phenocopying the NO-producing ones by reducing cell growth. The differences between CT26 and B16F10 at the basal level and IFN-γ-activation were initially investigated. At basal growth conditions, the B16F10 cells exhibited significantly lesser MHC class 1 surface expression and cell growth but higher lactate production than the CT26 cells (Fig.S8). Notably, IFN-γ-activated B16F10 cells exhibited much higher MHC class 1 induction regarding fold change (∼20-55-fold increase) than CT26 cells (∼2-5-fold increase). This observation indicated that the strength of response to IFN-γ in B16F10 cells is remarkably greater compared to CT26 cells (Fig.7A).

**Fig.6.**
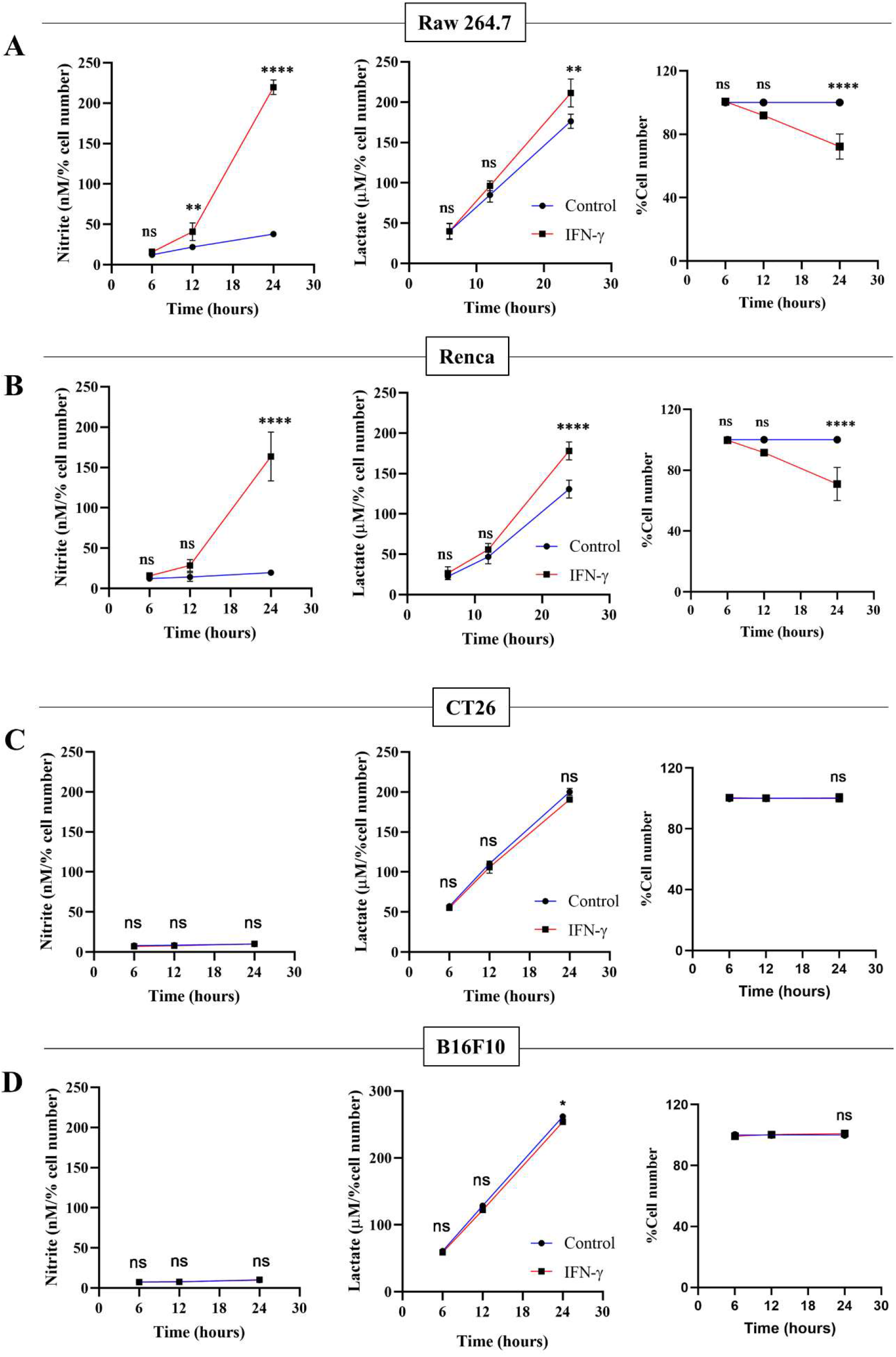
Only IFN-γ-activated NO-producing tumour cells increase lactate production and reduce cell number. Raw 264.7 (A), Renca (B), CT26 (C), and B16F10 (D) cell lines were treated with 10 U/mL IFN-γ in a 24-well plate. The cell culture medium was collected kinetically. The levels of nitrite and lactate were measured from the cell-free supernatant. The cell number was counted to derive the percent cell number. The statistical analyses were performed using two-way ANOVA with Tukey’s multiple comparisons tests for comparing nitrite and lactate levels and unpaired t-test for comparing the percent cell number. (ns), (*), (**), (***), (****) indicate non-significant difference and the statistical differences of p < 0.05, p < 0.01, p < 0.001, and p<0.0001 between the comparable indicated. Each data point is representative of the independent experiment. Data are represented as mean ± SD of 3-5 independent experiments.

**Fig.7.**
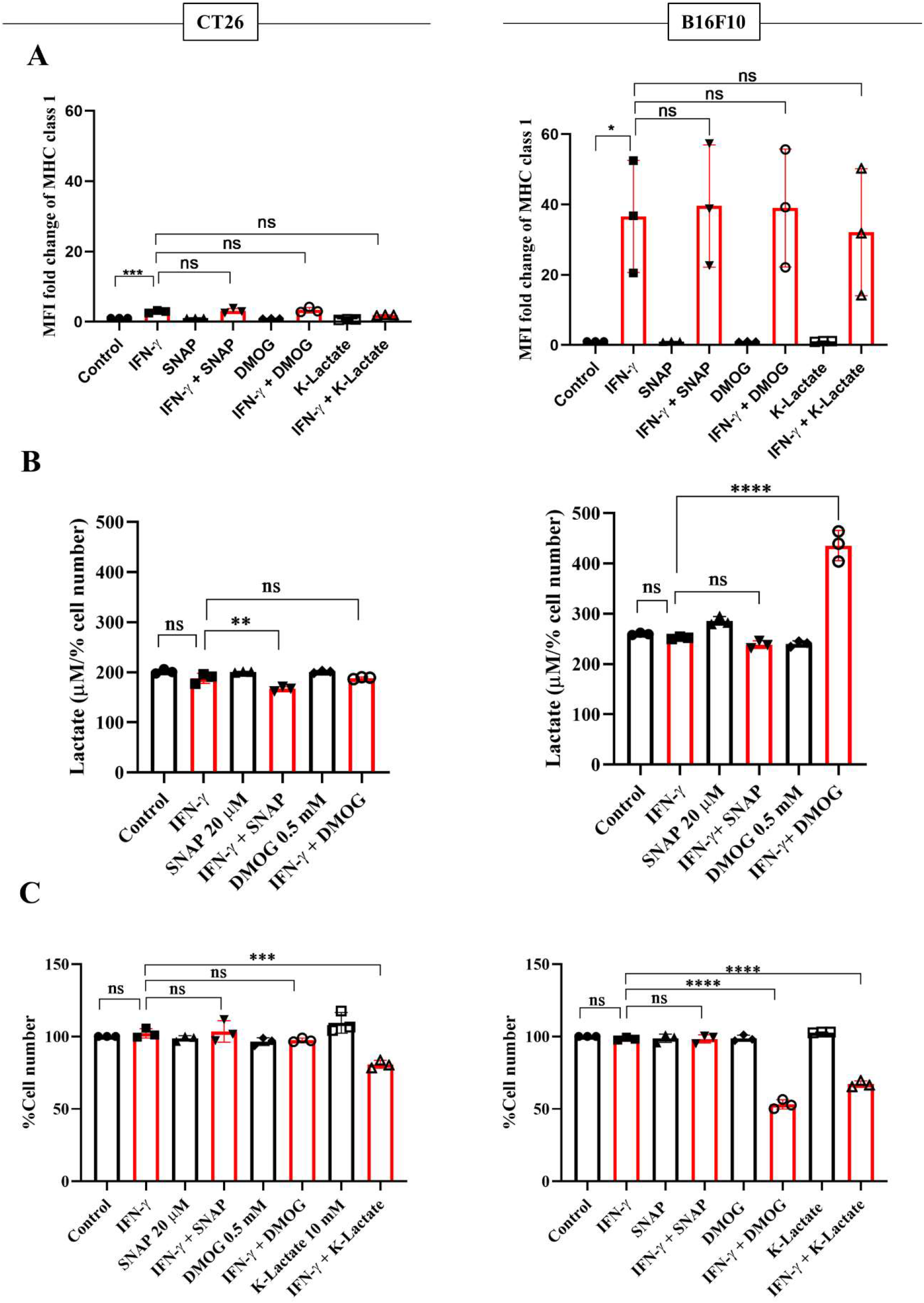
K-lactate in the presence of IFN-γ lowers the growth of non-NO-producing cell lines, CT26 and B16F10 cells. CT26 and B16F10 cells were treated with 10 U/ml IFN-γ and incubated alone or with pharmacological modulators (SNAP, an NO donor at 25 μM; DMOG, a stabilizer of HIF-1*α* at 0.5 μM and K-lactate as a metabolic reprogramming agent at 10 mM) for 24 hours. The cells were stained with an antibody to MHC class 1 to assess the surface expression of the MHC class 1 molecule (A). The level of lactate was measured from the cell-free supernatant. The lactate level was normalized to the percent cell number (B). The cell number was counted to derive the percent cell number (C). The statistical analyses were performed using ordinary one-way ANOVA with Sidak’s multiple comparisons tests. (ns), (*), (**), (***), (****) indicate non-significant difference and the statistical differences of p < 0.05, p < 0.01, p < 0.001, and p<0.0001 between the comparable indicated. Each data point is representative of the independent experiment. Data are represented as mean ± SD of 3 independent experiments.

NO, HIF-1α, and lactate were incorporated in CT26 and B16F10 cells using SNAP, DMOG, and potassium lactate. None of these pharmacological modulators affected IFN-γ-induced MHC class 1 surface expression in both CT26 and B16F10 cell lines (Fig. 7A). Apart from SNAP, none of the pharmacological modulators affected the NO amounts (Fig.S9B). Lactate levels and percent cell number remained unaffected upon adding SNAP in the presence of IFN-γ in B16F10 cells. DMOG did not affect the lactate levels and cell growth in CT26 cells upon IFN-γ-activation; however, it significantly increased lactate and reduced cell growth in B16F10 cells (Fig.7B, C). Most importantly, the exogenous addition of lactate into IFN-γ-activated CT26 and B16F10 cells significantly decreased the percent cell number, reducing cell growth (Fig.7C).

## Discussion

An optimal IFN-γ-activation is well-known to trigger anti-tumour responses in cancer cells by inhibiting angiogenesis, increasing Treg fragility, inducing tumour senescence, and triggering apoptosis and ferroptosis [18]. In this study, we demonstrated that IFN-γ-activated H6 cells underwent impairments in mitochondrial functions, glycolytic flux elevation, and extracellular acidification (Fig.1, S1). The primary modes of extracellular acidification are respiratory or metabolic. Enhancing mitochondrial function can release high levels of CO_2_ and contribute to respiratory acidification. However, efficient utilization of O_2_ under optimal mitochondrial membrane potential is necessary for CO_2_ release. Therefore, the reduced OCR upon IFN-γ-activation of H6 cells indicated that the IFN-γ-induced augmented glycolysis in H6 cells contributed to the extracellular acidification. The mitochondria of IFN-γ-activated H6 cells exhibited less mitochondrial membrane potential and oxygen consumption, indicating reduced dependency on mitochondrial oxidative phosphorylation for ATP production. The increased acidity can hamper enzyme functions and biochemical reactions. It is well-known that reducing the normal physiological pH by 0.1-0.2 in the human body leads to severe acidosis and can be fatal [42]. These observations indicated that IFN-γ-activated H6 cells phenocopy a similar metabolic rewiring, observed in activated macrophages to meet the cytostatic fate. The reduced tumour growth is less likely due to reduced pH because adding potassium lactate upon IFN-γ-activation of H6 cells reduced tumour growth without reducing pH (Fig.2F). These led us to investigate the detailed mechanism behind IFN-γ-induced glycolytic flux elevation.

NO derived from the non-malignant stromal components of tumours or NO-donor pharmacological agents often promotes tumour progression by angiogenesis, defective P53 functions, histone methylation, and metastasis [43]. However, intra-tumoral NO synthesis by NOS2 and peroxynitrite, a primary byproduct of excessive intracellular NO production, inhibits the mitochondrial Electron Transport System (ETS) complexes, derails the efficient electron transfer process to molecular oxygen, and produces mitochondrial superoxide radicals [44]. Superoxide radicals react with water molecules to produce hydrogen peroxide. Therefore, excessive NO production increases intracellular peroxynitrite and ROS levels. The elevated NO-induced nitrosative and oxidative damage triggers the caspase activation and apoptosis in NO-producing tumour cells [16]. This study led us to evaluate the IFN-γ response in tumour cells and categorize NO-producing tumour cells: H6, Raw264.7, and Renca, and non-NO-producing tumour cells: CT26 and B16F10. All IFN-γ-induced NO-producing cells produced heightened lactate amounts, accompanied with lowered cell growth, whereas non-NO-producing cells did not (Fig.1, 6). The IFN-γ-induced glycolytic flux enhancement in H6 cells depended on the inducible NO production (Fig.2B, C). Elevated ROS also contributed to heightened lactate production in these cells (Fig.2E). Furthermore, the increased glycolysis also promoted NO and ROS production in H6 cells, indicating the presence of reciprocal regulation between these processes (Fig.3A, B). Reciprocal regulation might strengthen the IFN-γ-signalling, culminating in apoptosis and cellular growth reduction.

Cellular oxygen sensing is tightly regulated to promptly adapt to hypoxic conditions by inducing glycolytic processes and meeting bioenergetic demands. Intracellular HIF-1*α* is hydroxylated by the Prolyl hydroxylases (PHDs) and degraded under normoxic conditions. On the other hand, low oxygen amounts in hypoxia limit oxygen availability for the PHD enzyme, leading to reduced hydroxylation of HIF-1*α* and its intracellular stabilization. HIF-1*α* is paramount in inducing glycolytic gene transcription and boosting glycolysis to produce ATP under hypoxia. However, HIF-1*α* can also be stabilized under immune non-hypoxic conditions by the influence of high amounts of NO and ROS [45]. IFN-γ-mediated inflammatory processes stabilize HIF-1*α* in a non-hypoxic manner. The IFN-γ-mediated HIF-1*α* stabilization is beneficial during Mycobacterial infections by inducing metabolic reprogramming toward glycolysis and bacterial clearance [26]. However, this regulation can be harmful during other conditions, such as the inflammation of the aortic valve [46]. Our study showed that IFN-γ-activated H6 cells transcriptionally upregulated *Hif1a* transcription and increased intracellular HIF-1*α* levels kinetically (Fig.4). The HIF-1*α* stabilization in H6 cells may have happened due to NO and ROS induction by IFN-γ. The HIF-1*α* might self-regulate its transcription to strengthen its actions. Alternatively, IFN-γ may induce the *Hif1a* transcription to increase intracellular HIF-1*α* levels and enhance glycolytic flux. IFN-γ reduced tumour growth by increasing NO and stabilizing HIF-1*α* to enhance glycolytic flux in H6 cells. The HIF-1*α*-induced glycolytic flux enhancement contributed to NO production, probably by supplying the substrate and various cofactors to NOS2 and contributing to ROS generation (Fig.5). Clinical datasets of inflammatory diseases revealed that the gene expression of HIF-1*α* and NOS2 are correlated in several diseases, like colon inflammation, Crohn’s disease, and mixed osteosarcoma (Fig. S5). These observations indicate that IFN-γ-induced *Nos2* and *Hif1a* transcription were correlated and may be functional in tissue inflammatory diseases.

In mammalian cells, lactate production increases during intense exercise and ischemia when the demand for ATP and oxygen exceeds the supply. Glucose-avid tumours produce lactate despite adequate oxygen tension: a phenomenon known as the Warburg effect. The build-up of lactate in stressed muscle, ischaemic tissues, and growing tumours has established lactate’s reputation as a harmful waste product [47,48]. Tumours also employ elevated lactate to promote an immunosuppressive state in the tumour microenvironment. The accumulation of lactate can induce the secretion of immunosuppressive factors to inhibit the immune response of NK cells and T cell [49]. However, our observation of heightened lactate production upon IFN-γ-activation of NO-producing tumour cells raised the question of whether lactate was a waste product. Or does lactate play an essential role in the IFN-γ signalling processes? Our experiments with the inhibition of lactate dehydrogenase and monocarboxylate transporters showed that IFN-γ-induced NO-depended augmentation of lactate production feeds into NO and ROS production processes. The IFN-γ-activated H6 cells may benefit from lactate shuttling in and out of the cells and supply metabolic intermediates through gluconeogenesis to support NO and ROS production. Ultimately, the reciprocally regulated circuitry lessens the tumoral growth. This speculation was further evidenced by using potassium lactate in IFN-γ-activated H6 cells, which increased NO production and decreased cell growth (Fig.3).

The interplay of NO, HIF-1*α*, and lactate in IFN-γ-activated H6 cells reducing tumour growth led us to investigate the effects of these factors in non-NO-producing tumour cells with IFN-γ: CT26 and B16F10. These tumour cell lines are commonly injected into mice, subcutaneously or orthotopically, to establish tumour growth [50,51]. Our investigations revealed that CT26 cells basally express higher MHC class 1 and grow faster compared to B16F10 cells. B16F10 cells were more robust producers of lactate than CT26 cells. IFN-γ-activation of CT26 cells did not increase MHC class 1 expression as robustly as B16F10 cells. Hence, B16F10 cells displayed more inducibility with IFN-γ activation, thereby increasing the response window compared to CT26 cells. This may be the reason why IFN-γ-activated B16F10 cells exhibited significant growth reduction when intracellular HIF-1*α* was pharmacologically stabilized. However, potassium lactate led to significant growth reduction upon IFN-γ-activation of both non-NO-producing cell lines, CT26 and B16F10 cells (Fig.7, S8, 9).

The implications of our findings will be discussed with respect to cancer immunotherapy. Despite considerable efforts, cancer immunotherapy shows limited success with immune checkpoint blockade, only ∼20.2% of patients achieved an objective response, and only ∼13% achieved multiyear durable responses [52]. IFN-γ influences all stages of tumour immunoediting and can have both pro-tumorigenic and anti-tumorigenic effects. At the early growth stage, IFN-γ eliminates the tumours by increasing the expression of MHC class 1, costimulatory molecules, the immunoproteasome, antigen processing and presentation, and reducing proliferation. However, sometimes poorly immunogenic and immune-evasive transformed cells establish equilibrium and escape immune-mediated killing to progress into cancer formation [53]. Patients with tumours that exhibit an active IFN-γ signature are more likely to respond favourably to immune checkpoint blockade [21,22]. Similarly, IFN-γ can modulate the expression of immunomodulatory molecules on tumour cells, making them more susceptible to CAR-T cell-mediated killing [20].

In conclusion, studying IFN-γ-driven immunometabolism is crucial for advancing cancer immunotherapy. Metabolic signatures associated with IFN-γ-driven tumour cell activation and function could serve as valuable indicators of immunotherapy efficacy. Our research enlightened the metabolic regulation of NO-producing H6 cells upon IFN-γ-activation. The IFN-γ-induced NO and ROS were important in elevating glycolytic flux and lactate production, possibly through HIF-1*α* stabilization. The HIF-1*α* function and glycolytic flux augmentation reciprocally regulated the NO and ROS production, strengthening the IFN-γ-activation signalling to induce nitrosative and oxidative stress and finally reducing tumour growth (Fig.8). Identifying patients more likely to respond to immunotherapy based on their responsiveness to IFN-γ, NO production and metabolic profiles can optimize treatment selection and minimize unnecessary exposure to ineffective therapies. Ultimately, we showed how non-NO-producing tumour cells upon IFN-γ-activation underwent cell growth reduction upon metabolic rewiring by potassium lactate. Therefore, our study shows that metabolic interventions like lactate or its analogues could make tumours more vulnerable to cancer immunotherapy.

**Fig.8. A.**
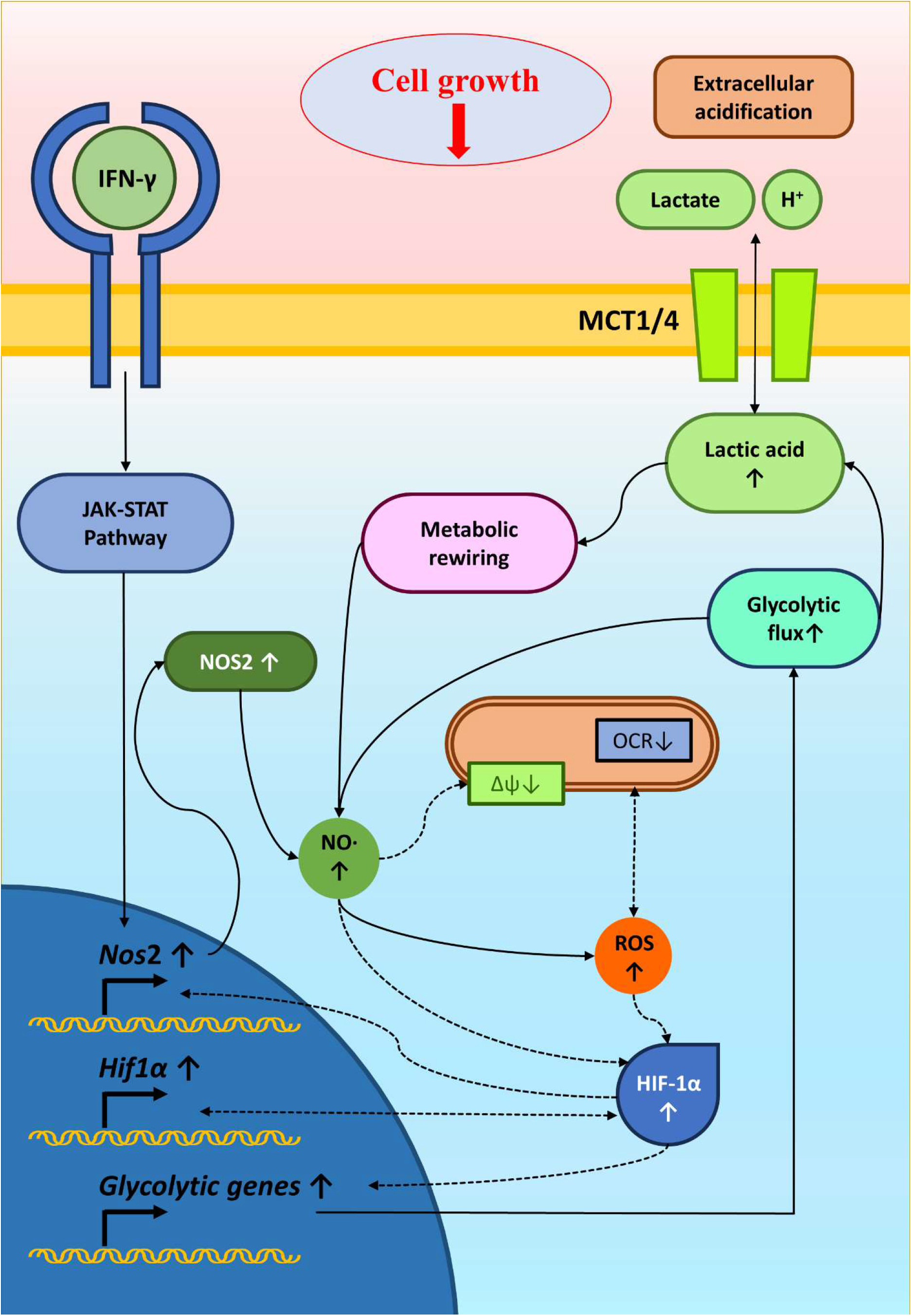
graphical illustration of IFN-γ-mediated enhanced NO and ROS increasing glycolysis and reducing tumour growth. IFN-γ-activation of H6 cells induces NO production and NO-mediated ROS generation. NO and ROS impair mitochondrial membrane potential and may reduce mitochondrial O_2_ consumption. NO, ROS and damaged mitochondrial function might stabilize HIF-1α levels. HIF-1α enhances the glycolytic flux upon IFN-γ-activation, possibly through increasing glycolytic gene expression. The enhanced glycolytic flux increases extracellular acidification and lactate accumulation. Furthermore, the heightened glycolysis and lactate reciprocally promoted the IFN-γ-induced NO and ROS production, reducing tumour cell growth. Solid arrows indicate connections with evidence. Dashed arrows indicate probable regulations.

## Conclusions

Investigating the glycolytic augmentation of IFN-γ-activated H6 cells provided essential information on the mechanism of tumoral nitrosative and oxidative signalling in reducing tumour growth. The major driving factors identified here: NO, HIF-1α, and lactate, played a significant role in IFN-γ-signalling-induced tumour growth reduction. Furthermore, incorporating these driving factors, especially potassium lactate in non-NO-producing tumour cells to reduce tumour growth, was paramount in showing the importance of reprogrammed glycolytic metabolism in IFN-γ-signalling. These studies reinforce the roles of metabolic interventions, which may improve cancer immunotherapy outcomes.

## Abbreviations

DMEM: Dulbecco’s modified minimal essential medium
ECAR: Extracellular Acidification Rate
HIF-1α: Hypoxia-inducible factor 1α
IFN-γ: Interferon-gamma
LNMA: NG-Methyl-L-arginine acetate salt
NO: Nitric Oxide
NOS: Nitric Oxide Synthase
OCR: Oxygen Consumption Rate

## Acknowledgments

We greatly appreciate the support from the members of the Divisional flow cytometry facility, IISc. This study was funded by SERB grant CRG/2021/004284, core grants from IISc and the DBT-IISc partnership program. In addition, the infrastructural support from DST-FIST to the Department of Biochemistry, IISc, is greatly appreciated. In addition, the support from all members of the laboratory is greatly appreciated.

## Author Contributions

AC designed experiments, performed, analyzed, interpreted data along with writing the paper. SJ performed RNA isolation and immunoblots from H6 cell line. AKK assisted in experiments and graphical illustrations. NSR assisted in cell culture experiments. AZ performed some experiments with glycolysis inhibitors. DN designed experiments, analyzed the data, revised, and approved the final manuscript.

