## Supplemental Information for "Interferon-γ lowers tumour growth by increasing glycolysis and lactate production in a nitric oxide-dependent manner: implications for cancer immunotherapy"

### Supplementary Information

#### Supplementary Table 1: Materials

| Reagents | | |
|---|---|---|
| **Name** | **Brand** | **Catalogue number** |
| 2-NBDG | Cayman chemicals (Ann Arbor, MI, United States) | 11046 |
| Glucose (HK) Assay Kit | Sigma-Aldrich (St. Louis, MO, USA) | GAHK-20 |
| Lactate Assay Kit (Colorimetric) | Cell Biolabs, Inc (San Diego, CA, USA) | MET-5012 |
| 24-well plate, flat bottom | Thermo Scientific (Roskilde, Denmark) | 142475 |
| 6-well plate, flat bottom | Thermo Scientific (Roskilde, Denmark) | 140675 |
| 96-well plate, flat bottom | Corning (Kennebunk, ME, USA) | 3599 |
| Immobilon Forte HRP substrate | EMD Millipore (Burlington, MA, USA) | WBULF0100 |
| Beta mercaptoethanol | Sigma-Aldrich (St. Louis, MO, USA) | M6250-100ML |
| Bicinchoninic Acid (BCA) Protein Assay | Gbioscience (St. Louis, MO, USA) | 786-570 |
| Chloroform | Sigma-Aldrich (St. Louis, MO, USA) | C2432-4X25ML |
| DCFDA | Sigma-Aldrich (St. Louis, MO, USA) | D6883 |
| TMRE | Thermofisher Scientific (St. Louis, MO, USA) | T669 |
| Diethyl Pyrocarbonate | Sigma-Aldrich (St. Louis, MO, USA) | D5758-25ML |
| Dulbecco's Modified Eagle Medium, High Glucose | Himedia (LBS Marg, Mumbai, India) | AT067-10X1L |
| Dulbecco's Phosphate Buffered Saline | Himedia (LBS Marg, Mumbai, India) | TS1006-10X1L |
| EDTA | Sigma-Aldrich (St. Louis, MO, USA) | E9884 |
| Ethanol | EMSURE® EMD Millipore (Billerica, MA, USA) | 654833 |
| Fetal Bovine Serum | Gibco (Fountain Park, Paisley, UK) | 10270106 |
| Gentamicin sulfate | Gbioscience (St. Louis, MO, USA) | AB1022 |
| HIF-1α antibody | Gbioscience (St. Louis, MO, USA) | ITA0611 |
| Interferon-γ | PeproTech (Rocky Hill, NJ, USA) | 315-05 |
| Isopropyl alcohol | Sigma-Aldrich (St. Louis, MO, USA) | I9030-100ML |
| L-Glutamine | Himedia (LBS Marg, Mumbai, India) | RM049-25G |
| Protease Phosphatase Arrest | Gbioscience (St. Louis, MO, USA) | 786-870 |
| β-actin antibody | Santa Cruz Biotechnology (Dallas, TX, USA) | sc-47778 |
| N-(1-Naphthyl) ethylenediamine dihydrochloride | Sigma-Aldrich (St. Louis, MO, USA) | N-5889 |
| ortho-Phosphoric acid | Sigma-Aldrich (St. Louis, MO, USA) | 612162 |
| Paraformaldehyde | Sigma-Aldrich (Bommasandra, Bangalore, India) | 158127 |
| PE-conjugated anti-mouse MHC Class I antibody | Ebioscience (San Diego, California, USA) | 12-5999-82 |
| Penicillin G Sodium Salt | Himedia (LBS Marg, Mumbai, India) | TC020 |

| | | |
|---|---|---|
| Seahorse XF calibrant | Agilent Seahorse XF (Santa Clara, CA, USA) | 100840-000 |
| XFp Glycolysis Stress Test Kit | Agilent Seahorse XF (Santa Clara, CA, USA) | 103017-100 |
| XFp Cell Mito Stress Test Kit | Agilent Seahorse XF (Santa Clara, CA, USA) | 103010-100 |
| Seahorse XFp FluxPak | Agilent Seahorse XF (Santa Clara, CA, USA) | 103022-100 |
| XF DMEM medium | Agilent Seahorse XF (Santa Clara, CA, USA) | 103575-100 |
| RevertAid First Strand cDNA Synthesis kit | Thermo Scientific (V.A.Graiciuno, Vilnius, Lithuania) | K1622 |
| Tris(hydroxymethyl)aminomethane | Sigma-Aldrich (St. Louis, MO, USA) | 252859 |
| Glycine | Sigma-Aldrich (St. Louis, MO, USA) | G8898 |
| SDS | Sigma-Aldrich (St. Louis, MO, USA) | L4390 |
| Tween 20 | Promega (Madison, USA) | H5151 |
| Methanol | Qualigens (Powai, Mumbai, India) | Q32407 |
| Sodium Azide | Himedia (LBS Marg, Mumbai, India) | RM123 |
| Sodium Bicarbonate | Sigma-Aldrich (St. Louis, MO, USA) | S5761-500g |
| Streptomycin | Gbioscience (St. Louis, MO, USA) | AB1037 |
| Sulfanilamide | Sigma-Aldrich (St. Louis, MO, USA) | S9251-100G |
| SYBR Green Mix | Gbioscience (St. Louis, MO, USA) | 786-5062 |
| HCl | SDFCL (Powai, Mumbai, India) | 20125L25 |
| TRI Reagent® | Sigma-Aldrich (St. Louis, MO, USA) | T9424-100ML |
| Trypan Blue solution | Sigma-Aldrich (St. Louis, MO, USA) | T6146-5G |
| Sodium chloride | Qualigens (Powai, Mumbai, India) | Q15915 |
| Skim milk | Himedia (Vadhani, Mumbai, India) | GRM1254 |
| PAGEmark™ Tricolor PLUS Protein Markers | Gbioscience (St. Louis, MO, USA) | 786-419 |
| Immobilon-P PVDF Membrane | Millipore (Peenya, Bangalore, India) | IPVH00010 |
| Trypsin-EDTA (0.25% Trypsin, 0.001% EDTA) | Himedia (LBS Marg, Mumbai, India) | TCL-014 |

**Instruments**

| Name | Brand |
|---|---|
| CFX Connect ™ | BioRad (Singapore) |
| FACSVerse ™ | BD Biosciences (USA) |
| Humidified CO2 Incubator | Sanyo (UK) |
| Infinite M200 Pro Multimode Microplate Reader | Tecan (Austria) |
| Nanodrop™ | Thermofisher Scientific (USA) |
| pH Meter LMPH-10 | Labman (India) |
| Thermomixer HM100 Pro | DLAB (India) |
| Seahorse XF analyzer (XF HS Mini) | Agilent technologies (USA) |
| Brightfield microscope | Euromex CCCD5 5.0 MP (India) |
| Semi-dry transfer apparaturs | GE Healthcare Biosciences Corp. (USA) |
| ChemiDoc Imaging System | BioRad (Singapore) |

**Supplementary Table 2: Details about the mouse adherent cell lines used in this study**

| Name | Mouse strain | Tissue origin | Cell type | Immortalization method | Antigen expression | Reference |
|---|---|---|---|---|---|---|
| H6 | A/J | Liver hepatocyte | Hepatoma | Information not available | $H\text{-}2^a$ | Thompson et al., 1981; Monaco et al., 1982 |
| Raw264.7 | BALB/c | Ascites | Macrophage | Abelson murine leukemia virus transformed | $H\text{-}2^d$ | Ralph and Nakoinz, 1977 |
| Renca | BALB/cCr | Renal cortex | Epithelial, adenocarcinoma | Arose spontaneously | $H\text{-}2^d$ | Murphy and Hrushesky, 1973 |
| CT26 | BALB/c | Large intestine | Colon Carcinoma | N-nitroso-N-methylurethane-(NNMU) induced | $H\text{-}2^d$ | Wang et al., 1995 |
| B16F10 | C57BL/6J | Skin | Melanoma | Arose spontaneously | $H\text{-}2^b$ | Lennicke et al., 2017 |

**Supplementary Table 3: Primers for RT-qPCR**

| Gene name | Forward Primer (5' - 3') | Reverse Primer |
|---|---|---|
| *Actb* | TGCTTCTAGGCGGACTGTTAC | TTTTGGGAGGGTGAGGGACTT |
| *Cd274* | CCCTTGACAGCTACTGCCTC | GGTTCGGCTATGCTCGTCTT |
| *Gapdh* | AGGTCGGTGTGAACGGATTTG | TGTAGACCATGTAGTTGAGGTCA |
| *Hif1α* | ATACCAACAGTAACCAACCT | GTCGACTGAGAAATGTCTTG |
| *Hk2* | GTGACTCTGATAGGTGCGAT | TAAGTCTCACTCCTGCCGA |
| *Irf1* | ATGCCAATCACTCGAATGCG | TTGTATCGGCCTGTGTGAATG |
| *Ldha* | AAGACAAACTCAAGGGCGAG | CTGGATTGGAGACGATCAGC |
| *Ldhb* | TGGTGGACAGTGCCTATGAA | TCTCAATGCCGTACATTCCC |
| *Nos2* | GTTCTCAGCCCAACAATACAAGA | GTGGACGGGTCGATGTCAC |
| *Vegfa* | CTATTCAGCGGACTCACCAG | GGGAGTGAAGAACCAACCTC |

**Supplementary Table 4. KEGG Pathway analysis of the HIF-1α-induced upregulated transcriptome**

| Pathway | Observed Gene count | Background Gene count | Matching Proteins in Network |
|---|---|---|---|
| HIF-1 signalling pathway | 13 | 111 | *Hk2, Eno2, Pfkl, Hif1a, Slc2a1, Egln1, Egln3, Pgk1, Aldoc, Ldha, Pfkfb3, Gapdh, Vegfa* |
| Glycolysis / Gluconeogenesis | 9 | 65 | *Hk2, Eno2, Pfkl, Pgk1, Pgm1, Aldoc, Ldha, Gapdh, Tpi1* |
| Fructose and mannose metabolism | 7 | 36 | *Hk2, Pfkl, Mpi, Pfkfb4, Aldoc, Pfkfb3, Tpi1* |
| Starch and sucrose metabolism | 5 | 31 | *Hk2, Gys1, Pygm, Pgm1, Gbe1* |
| Central carbon metabolism in cancer | 7 | 69 | *Hk2, Pfkl, Hif1a, Slc2a1, Slc16a3, Fgfr3, Ldha* |
| Biosynthesis of amino acids | 7 | 77 | *Eno2, Pfkl, Gpt2, Pgk1, Aldoc, Gapdh, Tpi1* |

### Supplementary figures:

A

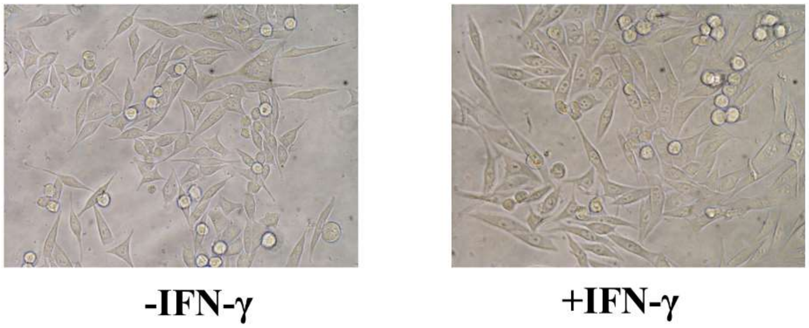

B

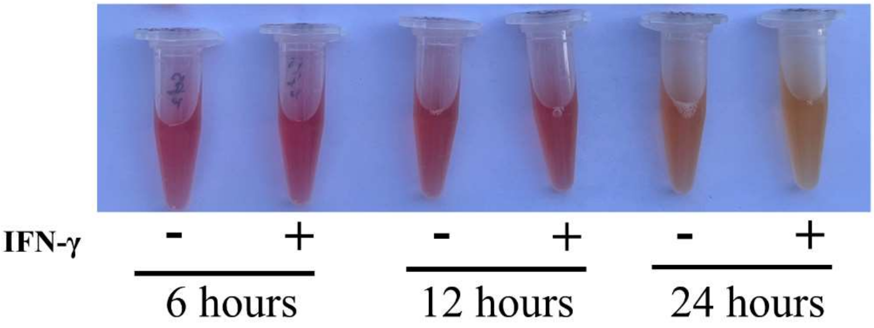

C

| Conditions | Control | IFN-γ |
|---|---|---|
| | 6.82 | 6.67 |
| | 6.85 | 6.6 |
| | 6.8 | 6.62 |
| | 6.87 | 6.69 |
| | 6.86 | 6.63 |
| **Mean** | **6.84** | **6.64** |
| **Standard Deviation** | **0.02** | **0.03** |

E

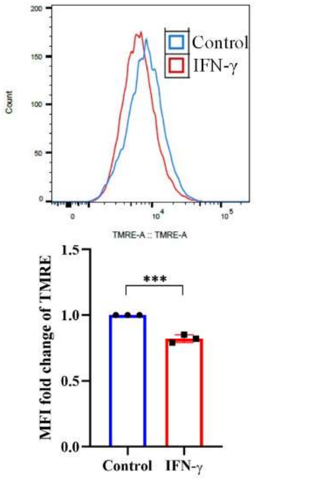

D

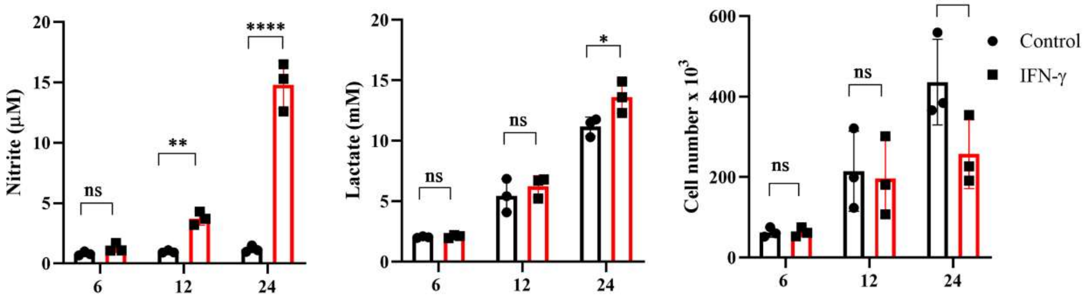

**Fig.S1. IFN-γ-activation of H6 cells increases NO, reduces cell number and extracellular pH without altering cytomorphology**

H6 hepatoma cells were treated with 10 U/mL IFN-γ for 24 hours. The cells were imaged using a 20X objective lens in a brightfield microscope (A). The cell-free supernatant was collected and photographed kinetically at the indicated time points (B). The pH of the cell-free supernatant was measured using a pH meter. Each unlabeled row represents an independent experiment. All independent experiments derived the mean and standard deviation (C). The nitrite and lactate concentration were measured in the cell-free supernatant. The total number of cells from 24-well plates after IFN-γ-activation is shown (D). Post-activation for 24 hours, the H6 cells were labeled with 10 μM TMRE dye for 15 minutes, and flow cytometric analysis was performed to compare mitochondrial transmembrane potential (E). The statistical analyses were performed using two-way ANOVA with Tukey's multiple comparisons tests (D) and unpaired t-test (E). (ns), (*), (**), (***), (****) indicate non-significant difference and the statistical differences of $p < 0.05$, $p < 0.01$, $p < 0.001$, and $p<0.0001$ between the comparable indicated. Each data point is representative of the independent experiment. Data are represented as mean ± SD of 3-5 independent experiments.

**A**

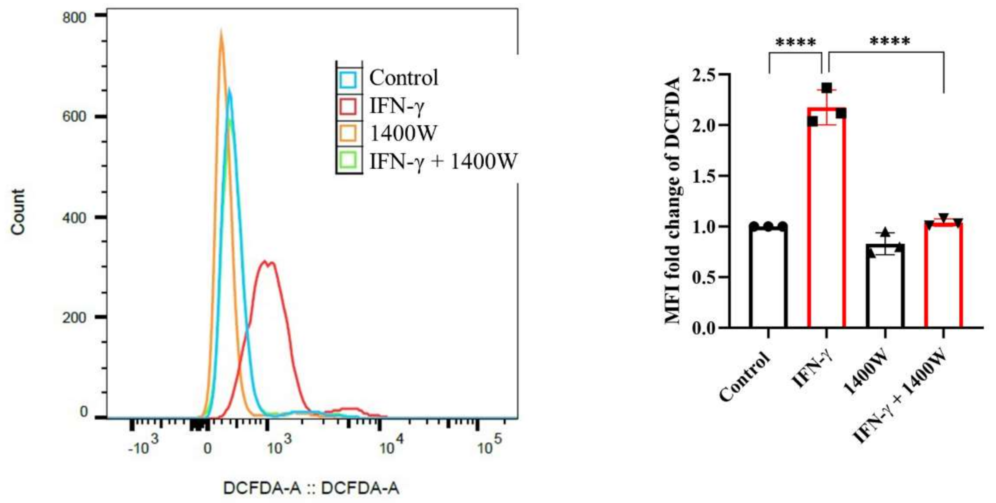
Control
IFN-γ
1400W
IFN-γ + 1400W
Count
DCFDA-A :: DCFDA-A
MFI fold change of DCFDA
****
****
Control
IFN-γ
1400W
IFN-γ + 1400W

**B**

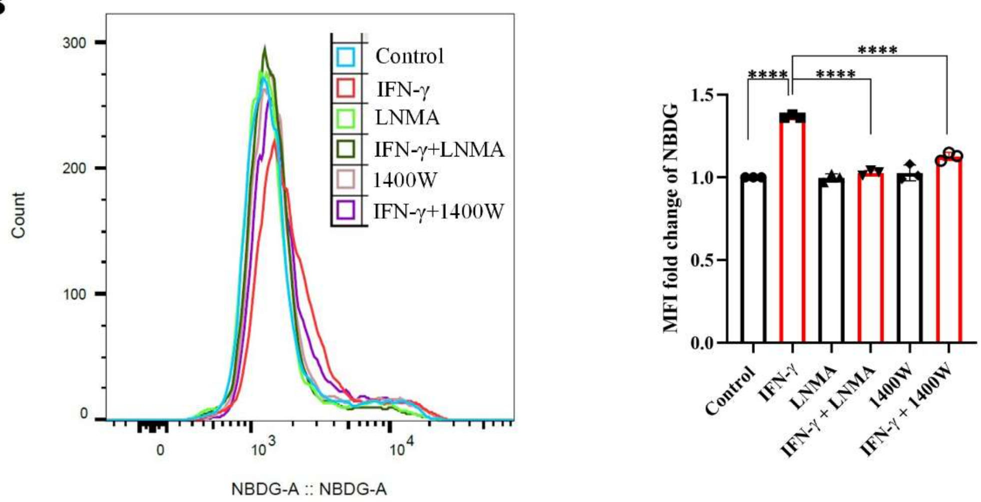
Control
IFN-γ
LNMA
IFN-γ+LNMA
1400W
IFN-γ+1400W
Count
NBDG-A :: NBDG-A
MFI fold change of NBDG
****
****
****
Control
IFN-γ
LNMA
IFN-γ + LNMA
1400W
IFN-γ + 1400W

**Fig.S2. IFN-γ-activated elevation of intracellular ROS and glucose uptake is NO-dependent.**

H6 cells were treated with 10 U/ml IFN-γ and incubated alone or with pharmacological inhibitors of NOS enzymes (LNMA at 200 uM and 1400W at 6 uM) for 24 hours to inhibit NO biosynthesis. The cells were incubated with 10 μM DCFDA or 100 μM 2-NBDG for 30 minutes, and flow cytometric analyses of intracellular ROS (A) or glucose uptake (B) were performed, respectively. The statistical analyses were performed using ordinary one-way ANOVA with Sidak's multiple comparisons tests. (ns), (*), (**), (***), (****) indicate non-significant difference and the statistical differences of $p < 0.05$, $p < 0.01$, $p < 0.001$, and $p<0.0001$ between the comparable indicated. Each data point is representative of the independent experiment. Data are represented as mean ± SD of 3 independent experiments.

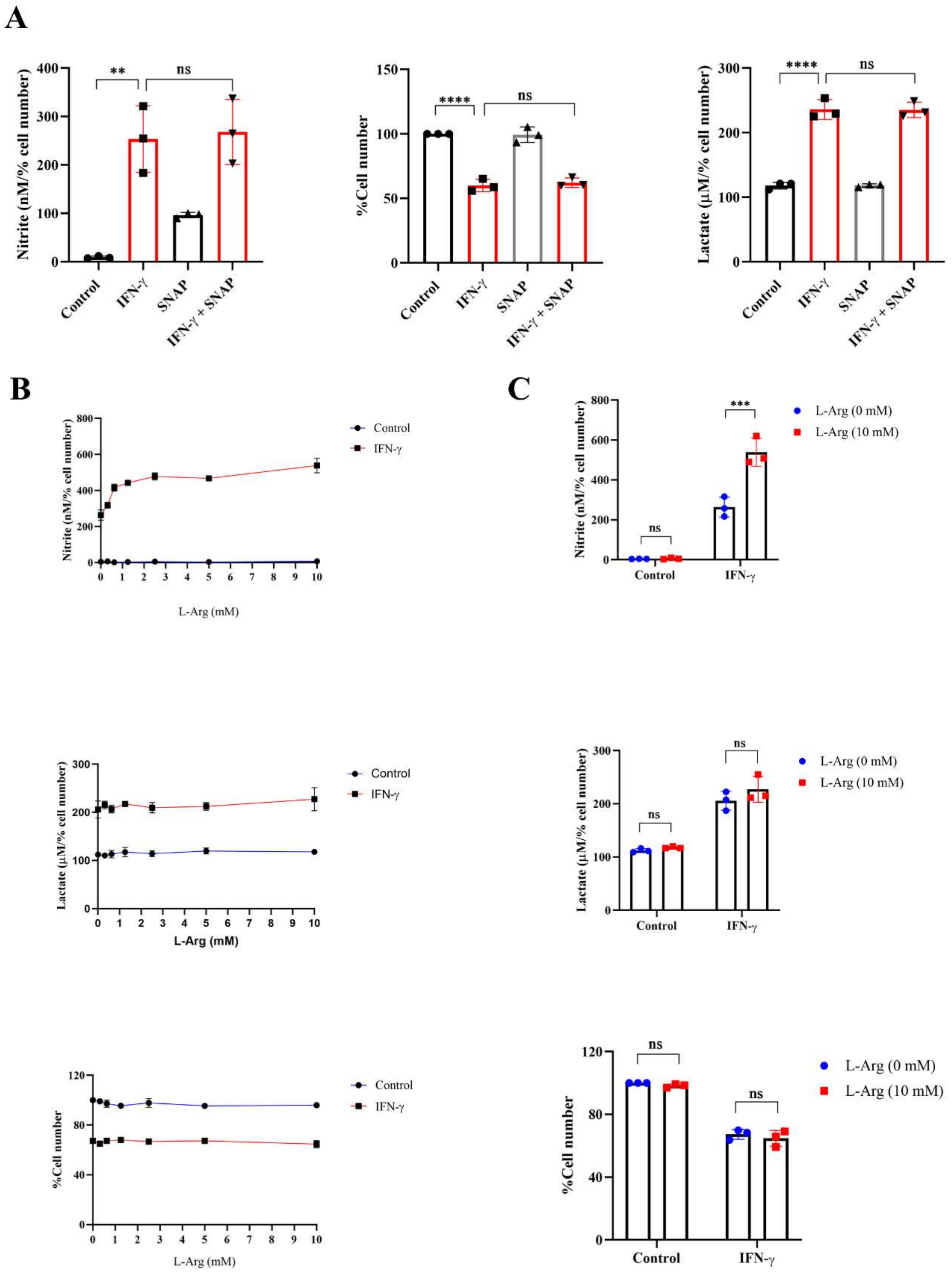
A
Nitrite (nM/% cell number)
%Cell number
Lactate (μM/% cell number)
**
****
ns
Control
IFN-γ
SNAP
IFN-γ + SNAP
B
C
Control
IFN-γ
L-Arg (mM)
L-Arg (0 mM)
L-Arg (10 mM)
***
ns

**Fig.S3. Exogenous L-arginine supplementation increases NO production in IFN-γ-activated H6 cells without affecting lactate production**

IFN-γ-activated H6 cells were treated with a NO-donor SNAP at 20 μM for 24 hours in a 24-well plate. The cell numbers were counted and represented as the percent cell number. The concentration of nitrite and lactate was estimated in the cell-free supernatant and normalized to the percent cell number (A). The IFN-γ-activated H6 cells were treated with L-arginine at the indicated concentrations for 24 hours in a 24-well plate. The cell numbers were counted and represented as the percent cell number. The nitrite and lactate concentrations were measured and normalized to the percent cell number (B, C). The statistical analyses were performed using ordinary one-way ANOVA with Sidak's multiple comparisons tests (A) and two-way ANOVA with Tukey's multiple comparisons tests (C). (ns), (*), (**), (***), (****) indicate non-significant difference and the statistical differences of $p < 0.05$, $p < 0.01$, $p < 0.001$, and $p<0.0001$ between the comparable indicated. Each data point is representative of the independent experiment. Data are represented as mean ± SD of 3-5 independent experiments.

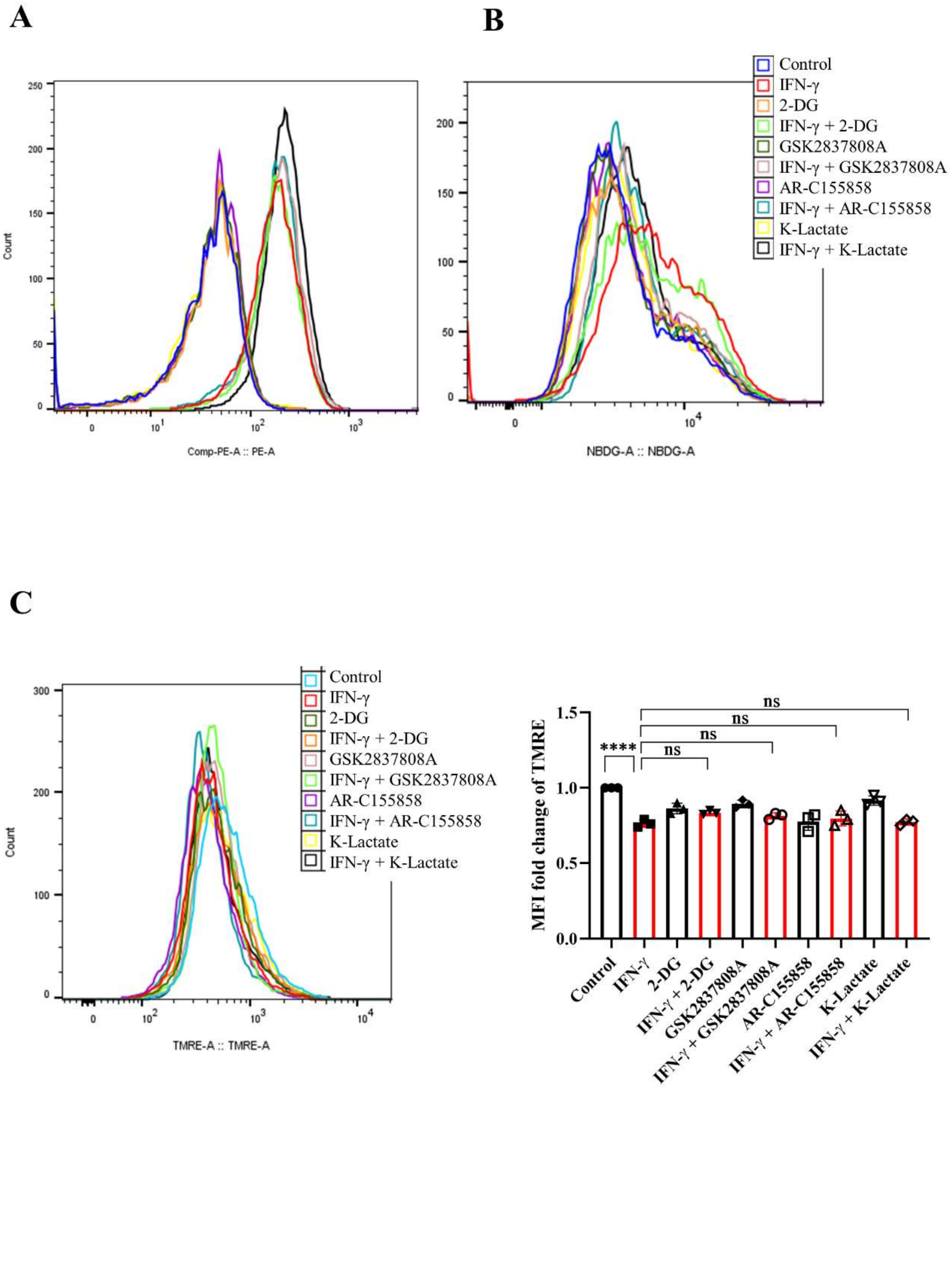
A
B
C
Count
Comp-PE-A :: PE-A
NBDG-A :: NBDG-A
TMRE-A :: TMRE-A
Control
IFN-γ
2-DG
IFN-γ + 2-DG
GSK2837808A
IFN-γ + GSK2837808A
AR-C155858
IFN-γ + AR-C155858
K-Lactate
IFN-γ + K-Lactate
MFI fold change of TMRE
****
ns

**Fig.S4. Glycolysis modulators do not affect the IFN-γ-mediated reduction of mitochondrial membrane potential**

H6 cells were treated with 10 U/ml IFN-γ and incubated alone or with pharmacological modulators of glycolysis (2-DG, a hexokinase inhibitor at 2.5 mM; GSK2837808A, a lactate dehydrogenase inhibitor at 125 nM; AR-C155858, a monocarboxylate transporter inhibitor at 6 nM and K-lactate as a metabolic reprogramming agent at 10 mM. The IFN-γ-activated H6 cells were assayed for glucose uptake and surface expression of MHC class 1 using flow cytometry in the presence of the glycolysis modulators. The cells were stained with an antibody to MHC class 1, and flow cytometric analysis of the surface expression of MHC Class 1 was performed (A). The cells were incubated with 100 μM 2-NBDG for 30 minutes, and flow cytometric analysis of glucose uptake was performed (B). Post-activation for 24 hours, the H6 cells were labeled with 10 μM TMRE dye for 15 minutes, and flow cytometric analysis was performed to compare mitochondrial transmembrane potential (C). The statistical analyses were performed using ordinary one-way ANOVA with Sidak's multiple comparisons tests (C). (ns), (*), (**), (***), (****) indicate non-significant difference and the statistical differences of $p < 0.05$, $p < 0.01$, $p < 0.001$, and $p<0.0001$ between the comparable indicated. Each data point is representative of the independent experiment. Data are represented as mean ± SD of 3 independent experiments.

A

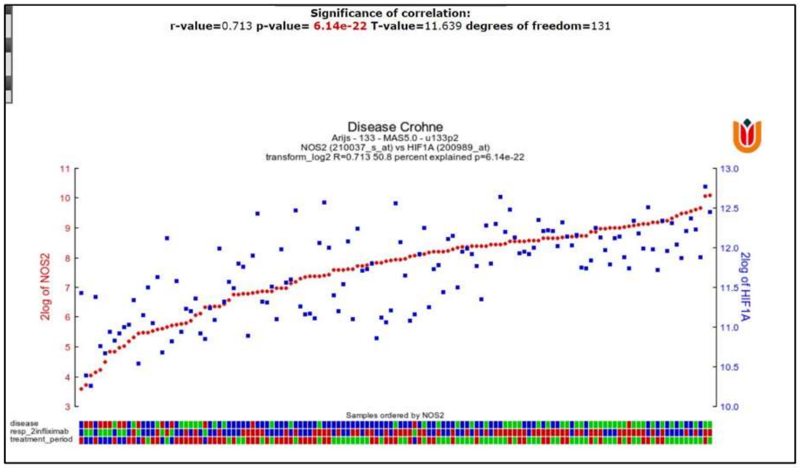
Significance of correlation:
r-value=0.713 p-value= 6.14e-22 T-value=11.639 degrees of freedom=131
Disease Crohne
Arijs - 133 - MAS5.0 - u133p2
NOS2 (210037_s_at) vs HIF1A (200989_at)
transform_log2 R=0.713 50.8 percent explained p=6.14e-22
2log of NOS2
2log of HIF1A
Samples ordered by NOS2
disease
resp_2infliximab
treatment_period

B

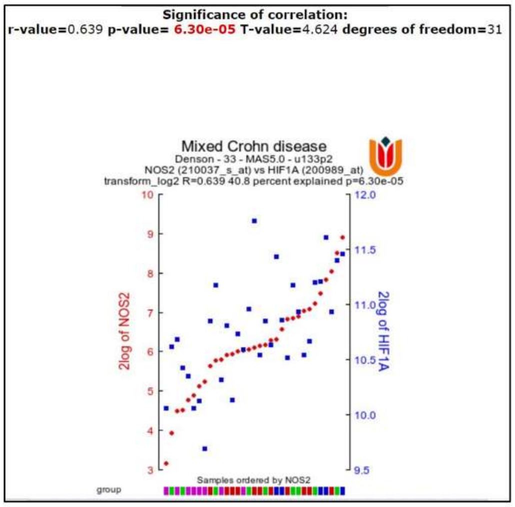
Significance of correlation:
r-value=0.639 p-value= 6.30e-05 T-value=4.624 degrees of freedom=31
Mixed Crohn disease
Denson - 33 - MAS5.0 - u133p2
NOS2 (210037_s_at) vs HIF1A (200989_at)
transform_log2 R=0.639 40.8 percent explained p=6.30e-05
2log of NOS2
2log of HIF1A
Samples ordered by NOS2
group

C

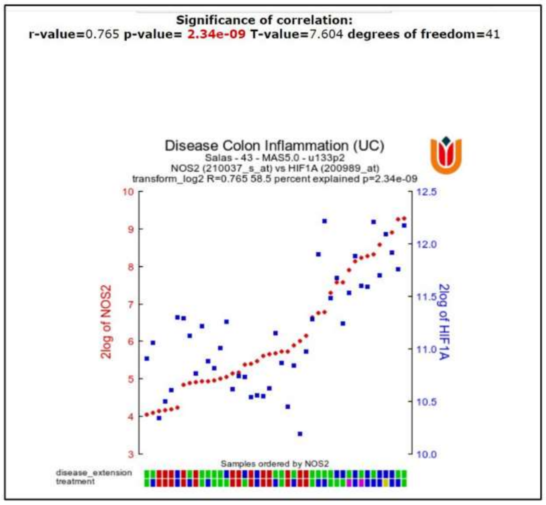
Significance of correlation:
r-value=0.765 p-value= 2.34e-09 T-value=7.604 degrees of freedom=41
Disease Colon Inflammation (UC)
Salas - 43 - MAS5.0 - u133p2
NOS2 (210037_s_at) vs HIF1A (200989_at)
transform_log2 R=0.765 58.5 percent explained p=2.34e-09
2log of NOS2
2log of HIF1A
Samples ordered by NOS2
disease_extension
treatment

D

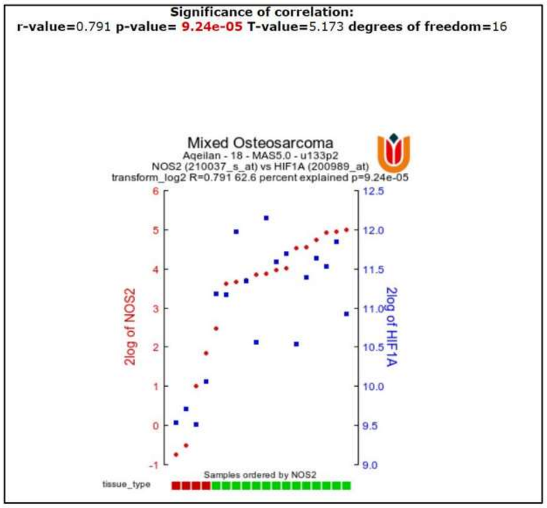
Significance of correlation:
r-value=0.791 p-value= 9.24e-05 T-value=5.173 degrees of freedom=16
Mixed Osteosarcoma
Aqeilan - 18 - MAS5.0 - u133p2
NOS2 (210037_s_at) vs HIF1A (200989_at)
transform_log2 R=0.791 62.6 percent explained p=9.24e-05
2log of NOS2
2log of HIF1A
Samples ordered by NOS2
tissue_type

**Fig.S5. The expression of NOS2 and HIF1A is correlated in clinical datasets of colon inflammation and osteosarcoma**

The R2 genomics analysis and visualization platform (R2 Genomics Analysis and Visualization Platform (amc.nl)) were used to investigate the correlative expression of NOS2 and HIF1A in the disease datasets. The results from Crohn's disease (A, B), colon inflammation (C), and mixed osteosarcoma (D) are represented.

**A**

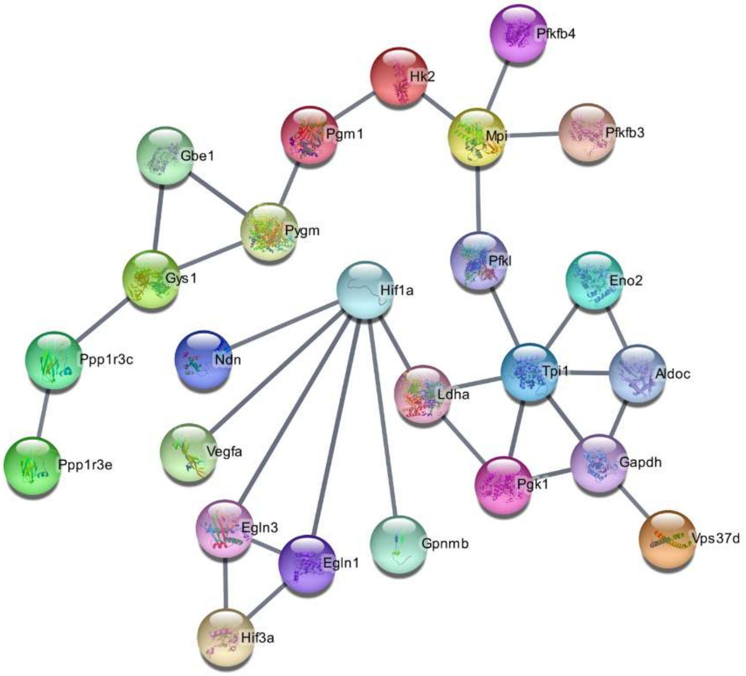
Pfkfb4
Hk2
Pgm1
Mpi
Pfkfb3
Gbe1
Pygm
Pfkl
Gys1
Hif1a
Eno2
Ppp1r3c
Ndn
Tpi1
Aldoc
Ldha
Ppp1r3e
Vegfa
Gapdh
Pgk1
Egln3
Vps37d
Gpnmb
Egln1
Hif3a

**B**

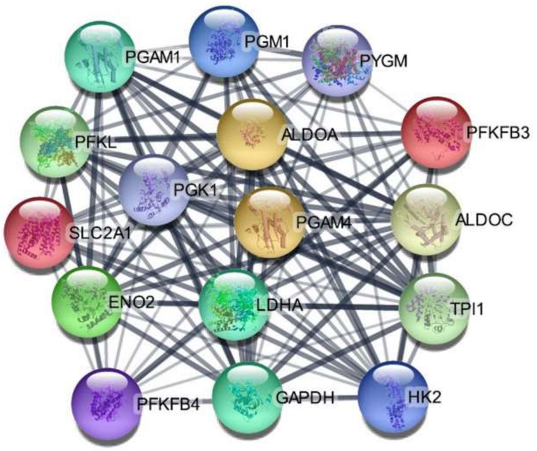
PGAM1
PGM1
PYGM
PFKL
ALDOA
PFKFB3
PGK1
SLC2A1
PGAM4
ALDOC
ENO2
LDHA
TPI1
PFKFB4
GAPDH
HK2

**C**

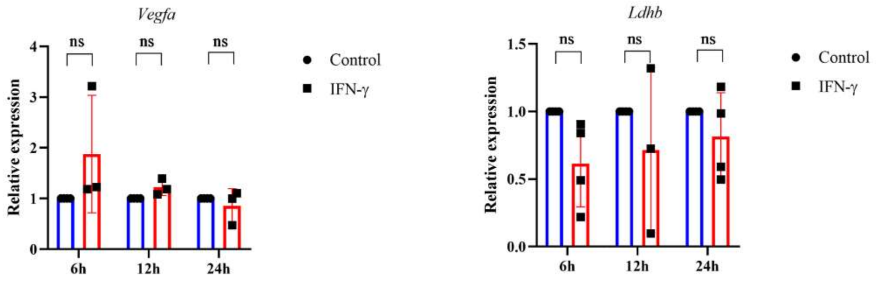
Vegfa
ns
Control
IFN-γ
Relative expression
0
1
2
3
4
6h
12h
24h
Ldhb
0.0
0.5
1.0
1.5

**Fig.S6. Protein-protein interaction network from human Hif1a-induced gene signature demonstrates interconnected clusters of glycolytic genes**

A protein-protein interaction network of human Hif1a-induced differentially expressed genes was built in String, and the topmost interconnected cluster was identified using the k-means clustering method (A). The MCODE algorithm in Cytoscape identified the metabolic gene cluster in the network of HIF-1a-induced gene expression (B). H6 cells were treated with 10 U/ml IFN-γ in a 6-well plate for the indicated time, and total RNA was extracted. RT-qPCR was performed to quantify the relative expression of *Vegfa* and *Ldhb* (C). The statistical analyses were performed using two-way ANOVA with Tukey's multiple comparisons tests (C). (ns), (*), (**), (***), (****) indicate non-significant difference and the statistical differences of $p < 0.05$, $p < 0.01$, $p < 0.001$, and $p<0.0001$ between the comparable indicated. Each data point is representative of the independent experiment. Data are represented as mean ± SD of 2-4 independent experiments.

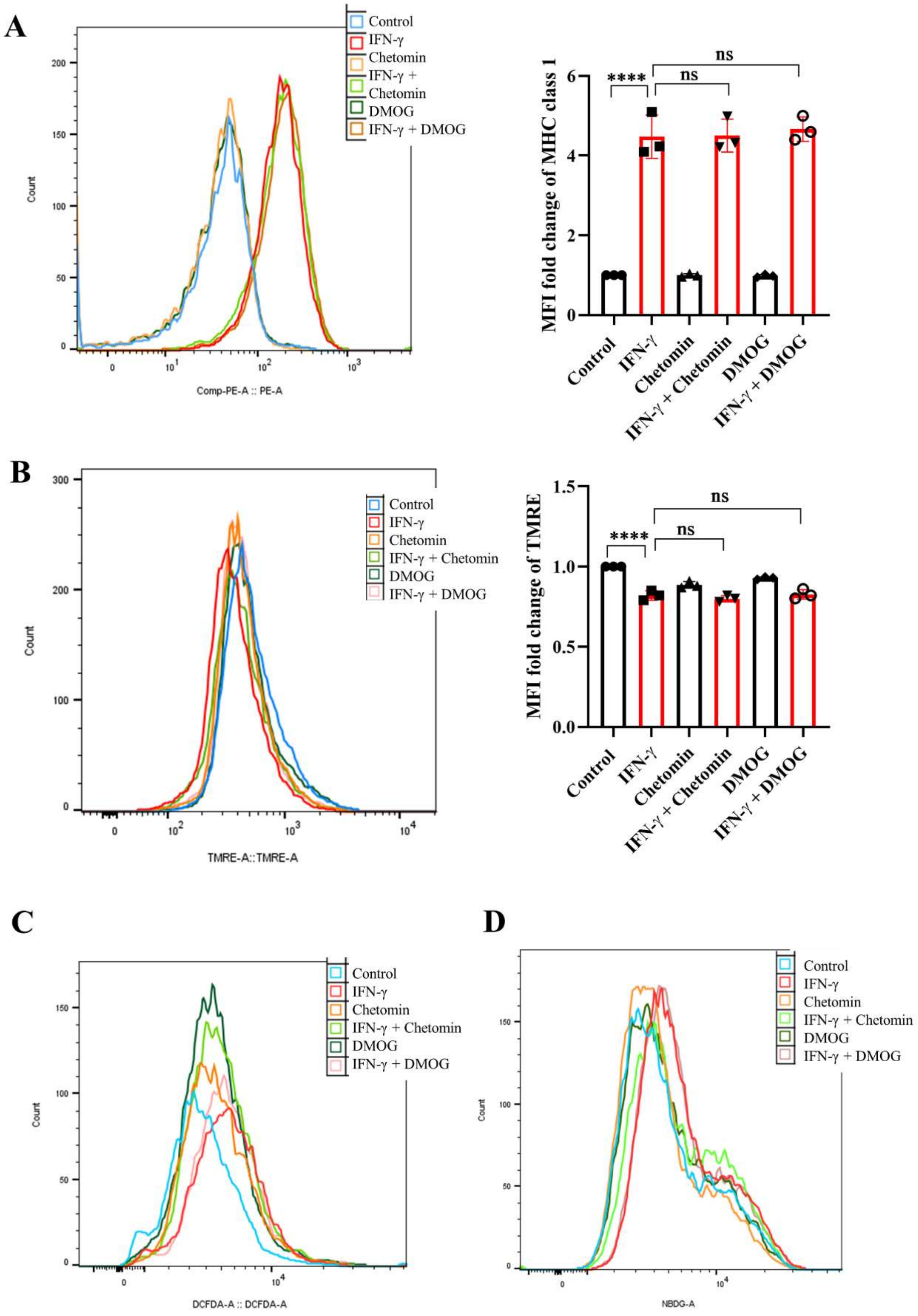
A
Control
IFN-γ
Chetomin
IFN-γ + Chetomin
DMOG
IFN-γ + DMOG
Count
Comp-PE-A :: PE-A
MFI fold change of MHC class 1
****
ns
ns
Control
IFN-γ
Chetomin
IFN-γ + Chetomin
DMOG
IFN-γ + DMOG
B
Control
IFN-γ
Chetomin
IFN-γ + Chetomin
DMOG
IFN-γ + DMOG
Count
TMRE-A::TMRE-A
MFI fold change of TMRE
****
ns
ns
Control
IFN-γ
Chetomin
IFN-γ + Chetomin
DMOG
IFN-γ + DMOG
C
Control
IFN-γ
Chetomin
IFN-γ + Chetomin
DMOG
IFN-γ + DMOG
Count
DCFDA-A :: DCFDA-A
D
Control
IFN-γ
Chetomin
IFN-γ + Chetomin
DMOG
IFN-γ + DMOG
Count
NBDG-A

**Fig.S7. Modulating HIF1A function does not affect IFN-γ-induced MHC class 1 surface expression elevation and lowering of mitochondrial membrane potential.**

H6 cells were treated with 10 U/ml IFN-γ for 24 hours. The IFN-γ-activated cells were incubated alone or with pharmacological modulators of HIF1A function (Chetomin, an inhibitor of HIF1A at 12.5 nM, and DMOG, a stabilizer of HIF1A at 0.5 uM). The cells were stained with an antibody to MHC class 1, and flow cytometric analysis of the surface expression of MHC Class 1 was performed (A). The cells were labeled with 10 μM TMRE dye for 15 minutes, and flow cytometric analysis was performed to compare mitochondrial transmembrane potential (B). The cells were incubated with 10 μM DCFDA for 30 minutes, and flow cytometric analysis of intracellular ROS was performed (C). The cells were incubated with 100 μM 2-NBDG for 30 minutes, and flow cytometric analysis of glucose uptake was performed (D). The statistical analyses were performed using ordinary one-way ANOVA with Sidak's multiple comparisons tests (C). (ns), (*), (**), (***), (****) indicate non-significant difference and the statistical differences of $p < 0.05$, $p < 0.01$, $p < 0.001$, and $p<0.0001$ between the comparable indicated. Each data point is representative of the independent experiment. Data are represented as mean ± SD of 3 independent experiments.

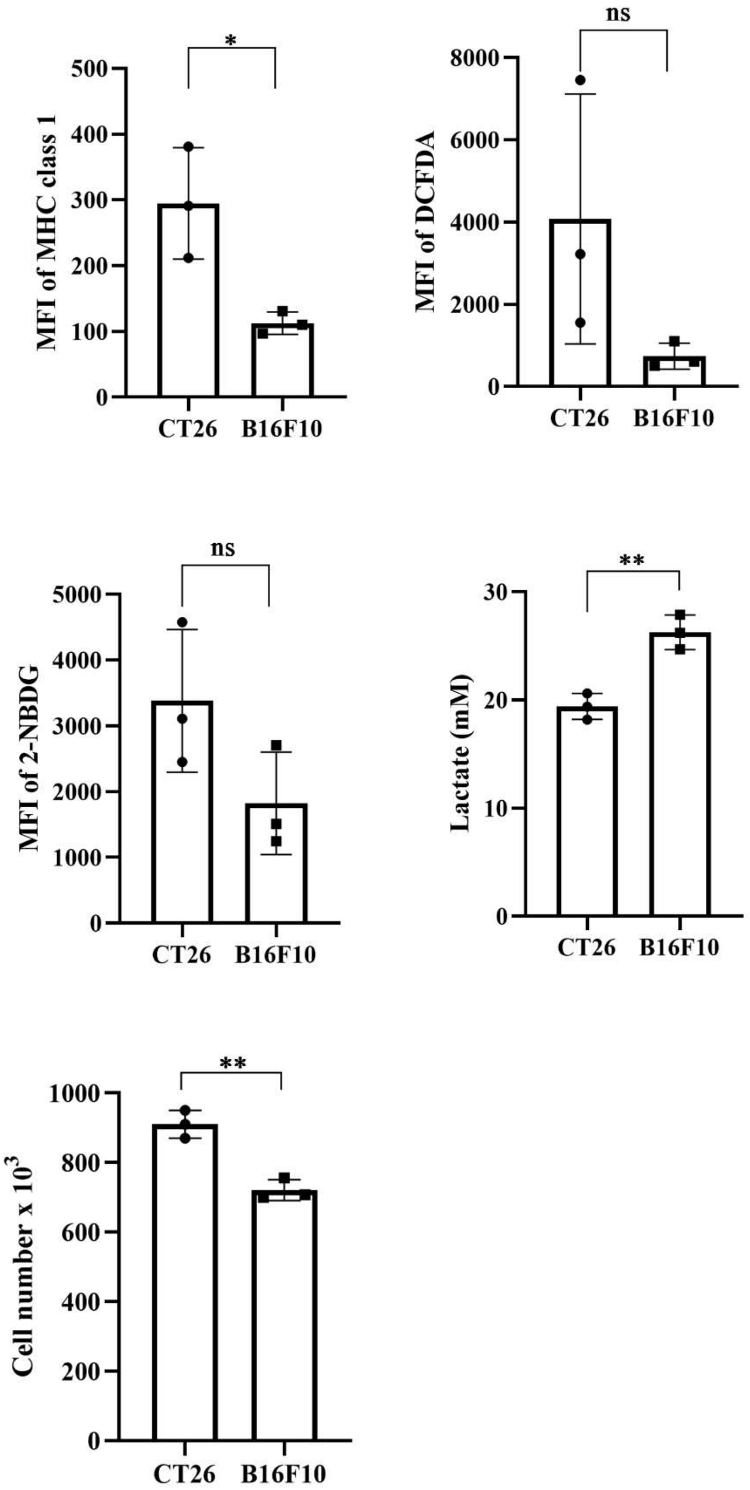
*
MFI of MHC class 1
500
400
300
200
100
0
CT26
B16F10
ns
MFI of DCFDA
8000
6000
4000
2000
0
CT26
B16F10
ns
MFI of 2-NBDG
5000
4000
3000
2000
1000
0
CT26
B16F10
**
Lactate (mM)
30
20
10
0
CT26
B16F10
**
Cell number x $10^3$
1000
800
600
400
200
0
CT26
B16F10

**Fig.S8. B16F10 cells express lower amounts of MHC class 1 and produce more lactate compared to CT26 cells**

CT26 and B16F10 cells were seeded in a 24-well plate at a density of 0.05 million cells per well and incubated for 24 hours. The cells were stained with an antibody to MHC class 1 to assess the surface expression of the MHC class 1 molecule. The cells were incubated with 10 μM DCFDA for 30 minutes, and flow cytometric analysis of intracellular ROS was performed. The cells were incubated with 100 2-NBDG for 30 minutes, and flow cytometric analysis of glucose uptake was performed. The lactate amounts were measured from the cell-free supernatant. The cell numbers were counted to derive each well's total number of cells. The statistical analyses were performed using the unpaired t-test. (ns), (*), (**), (***), (****) indicate non-significant difference and the statistical differences of $p < 0.05$, $p < 0.01$, $p < 0.001$, and $p<0.0001$ between the comparable indicated. Each data point is representative of the independent experiment. Data are represented as mean ± SD of 3 independent experiments.

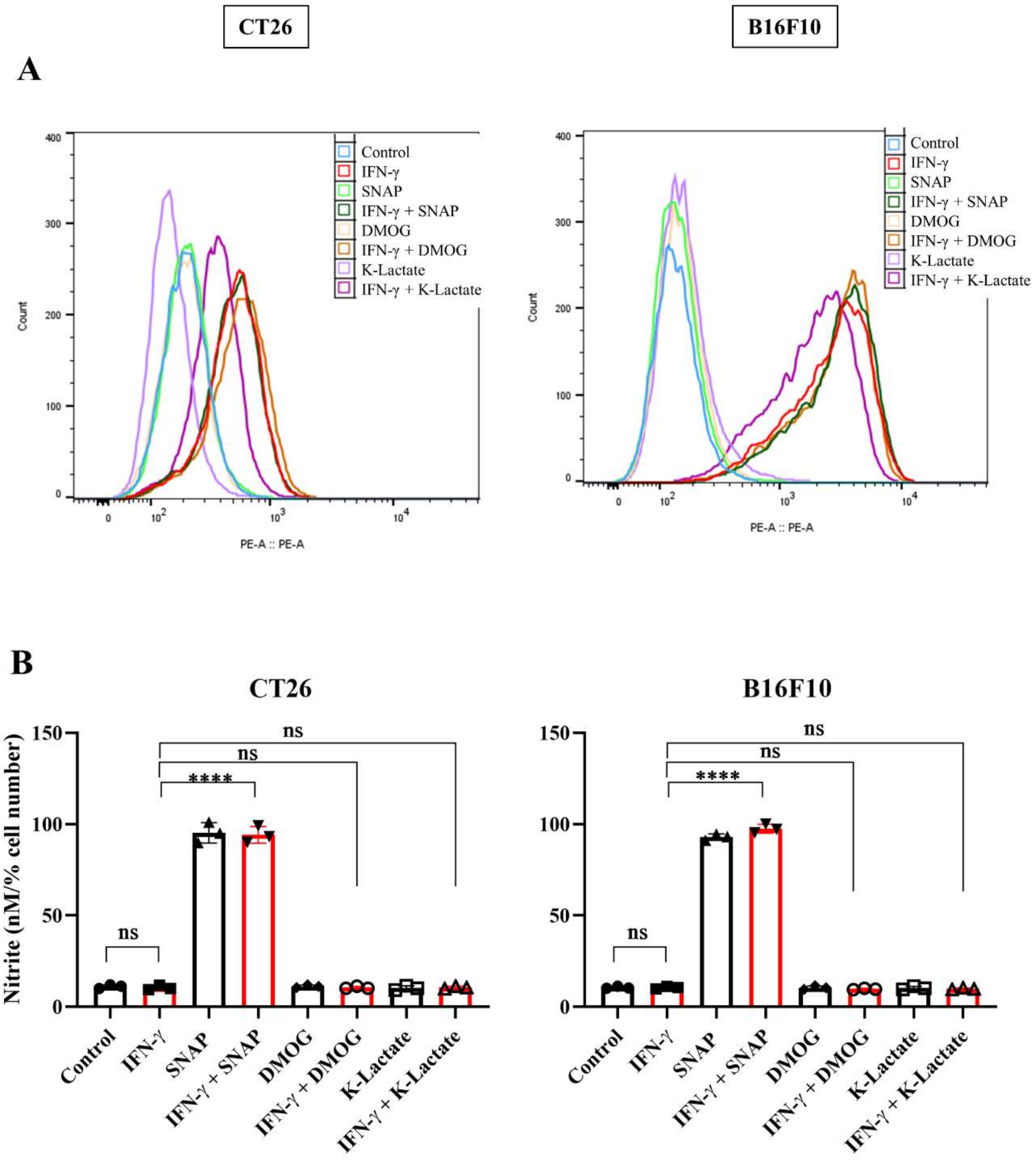
CT26
B16F10
A
Control
IFN-γ
SNAP
IFN-γ + SNAP
DMOG
IFN-γ + DMOG
K-Lactate
IFN-γ + K-Lactate
Count
PE-A :: PE-A
B
CT26
B16F10
Nitrite (nM/% cell number)
ns
****
Control
IFN-γ
SNAP
IFN-γ + SNAP
DMOG
IFN-γ + DMOG
K-Lactate
IFN-γ + K-Lactate

**Fig.S9. DMOG and K-lactate did not affect the NO amounts in CT26 and B16F10 cells**

CT26 and B16F10 cells were seeded in a 24-well plate at a density of 0.05 million cells per well and incubated for 24 hours. The cells were stained with an antibody to MHC class 1 to assess the surface expression of the MHC class 1 molecule (A). The cell number was counted to derive each well's total number of cells. The nitrite amounts were measured from the cell-free supernatant and normalized to the percent cell number (B). The statistical analyses were performed using the ordinary one-way ANOVA with Sidak's multiple comparisons tests (B). (ns), (*), (**), (***), (****) indicate non-significant difference and the statistical differences of $p < 0.05$, $p < 0.01$, $p < 0.001$, and $p<0.0001$ between the comparable indicated. Each data point is representative of the independent experiment. Data are represented as mean ± SD of 3 independent experiments.

### Supplementary methods

#### Cell number and FCCP optimization for Seahorse XF analyses

H6 cells were seeded at the density of 4000 (Well number: B, C, D) and 8000 (Well number: E, F, G) per well in triplicates. The cells were adhered for 8 h, washed to remove floating cells and incubated in the $CO_2$ incubator for 24 h. Subsequently, the cells were treated with the supplemented seahorse assay medium, as described for mitostress test in the manufacturer's protocol. A, B, C & D wells had FCCP as 0 µM in port B, 0.125 µM in port C, and 0.25 µM in port D. E, F, G, & H wells had 0.5 µM in port B, 1 µM in port C, and 2 µM in port D. Oligomycin was added in port A of all wells. According to the results, the cell number of 8000 per well and FCCP concentration of 1 µM were finalized for the seahorse XF analyses.

#### Reconstitution of the oligonucleotide primers

The lyophilized Oligonucleotide vials were centrifuged at 15000 g for 10 minutes. To prepare a 100 µM stock, nuclease-free water was added at the recommended volume provided on the technical datasheet. The vials were gently tapped and centrifuged at 130 g (pulse). The vials were kept at 56°C for 20 min and agitated intermittently after every 5 min. A pulse of 13,300 g was given 1-2 times. A 10 µM working solution was prepared in nuclease-free microfuge tubes. The primers were stored at -20°C till further usage.

#### Brightfield microscopy

~$10^4$ H6 cells were seeded on 96-well plates and treated with or without IFN-γ. Images were captured after 24 hours of treatment under a 40X objective lens, fitted in a brightfield microscope.
